# Genome-wide in vivo screen of circulating tumor cells identifies SLIT2 as a regulator of metastasis

**DOI:** 10.1101/2021.08.20.457126

**Authors:** Fan Xia, Yuan Ma, Kangfu Chen, Bill Duong, Sharif Ahmed, Randy Atwal, David Philpott, Troy Ketela, Jennifer Pantea, Sichun Lin, Stephane Angers, Shana O. Kelley

## Abstract

Circulating tumor cells (CTCs) break free from primary tumors and travel through the bloodstream and lymphatic system to seed metastatic tumors, which are the major cause of death from cancer. The identification of the major genetic factors that enhance production and persistence of CTCs in the bloodstream at a whole genome level would enable more comprehensive molecular mechanisms of metastasis to be elucidated and the identification of novel therapeutic targets, but this remains a challenging task due to the heterogeneity and extreme rarity of CTCs. Here, we describe the first *in vivo* genome-wide CRISPR KO screen using CTCs directly isolated from a mouse xenograft. This screen elucidated *SLIT2* – a gene encoding a secreted protein acting as a cellular migration cue – as the most significantly represented gene knockout in the CTC population. *SLIT2* knockout cells are highly metastatic with hypermigratory and mesenchymal phenotype. Reduced expression of *SLIT2* is observed in human tumors, indicating its role as a negative modulator of tumor progression and metastasis.

## Main

Metastasis accounts for over 90% deaths from cancer, yet remains poorly understood and largely incurable^1–3^. Despite advances in cancer treatment that significantly reduce morbidity and mortality of various cancer types, metastasis still leads to poor prognosis of cancer patients and treating metastasis remains a tremendous challenge ^4,5^ In order to develop more effective therapies and improve patient outcomes, further progress elucidating the fundamental biology of metastasis is critical to reveal mechanisms at the molecular level and provide insights regarding new therapeutic targets. Comparing the genetic background between primary tumors and metastatic lesions using whole exome sequencing (WES) and whole genome analyses has facilitated the identification of genetic factors that drive tumor progression and dissemination^6–8^. However, deciphering drivers of metastasis solely based on the genetic information from solid tumors is limiting due to genetic divergence and tumor heterogeneity^7–9^.

As circulating tumor cells (CTCs) leave primary tumors and seed metastatic lesions, molecular characterization of CTCs is critical to facilitate comprehensive understanding of the metastatic processes. Progress in the area of CTC enrichment and single-cell sequencing technologies has enabled identification of CTC-specific mutations in cancer patients at the genomic level^10–12^. However, efficient capture of CTCs and their unbiased genomic amplification for WES are still challenging due to the rarity and fragility of CTCs^11–15^. In addition, particular mutations observed in individual patient CTCs could be anecdotal, with low applicability and weak prognostic potential for large patient populations. Thus, new technologies and platforms are needed to comprehensively and systematically study the genetic factors underlying the CTC phenotypes that contribute to the metastatic process.

Here, we report a genome-wide CRISPR knockout (KO) screen designed to identify genes contributing to CTC dissemination *in vivo*. Xenografted tumors were seeded with pools of CRISPR-edited cells with each cell possessing loss-of-function of one gene in the human genome. Using a high-performance approach for CTC capture directly from blood coupled with next generation sequencing (NGS) of barcoded sgRNAs, gene knockouts that promoted CTC abundance were identified. This approach allowed the systematic identification of possible metastasis-promoting genetic factors at the CTC level with minimal bias. Previous genome-wide *in vivo* CRISPR screens identifying metastatic factors primarily used solid tumors or CTCs cultured *ex vivo* ^16,17^. Our approach however, allowed genome-wide *in vivo* CRISPR screening to be performed on fresh CTCs directly collected from liquid biopsy without extensive culturing. Avoiding *ex vivo* culturing of CTCs is important to prevent possible changes in morphology and proliferation of CTC subpopulations ^18,19^, which could affect representations of subsets of sgRNAs in the library. Our *in vivo* CTC CRISPR KO screen elucidated SLIT2 as a secreted factor that promoted the epithelial-to-mesenchymal transition (EMT) of tumor cells, resulting in enhanced intravasation. *SLIT2* KO cells had increased expression of complex I and were hypersensitive to the treatment of complex I inhibitor rotenone. Loss of *SLIT2* increased CTC numbers and the extent of metastasis of prostate cancer in an animal model.

## Results

### Genome-wide in vivo CRISPR-Cas9 knockout screen on CTCs

Genome-wide CRISPR screens are powerful tools for the systematic and unbiased identification of genes involved in metastasis^16,17,20^. Combining a CRISPR-Cas9 genome-wide editing system (TKOv3: Toronto KnockOut version 3 CRISPR library)^21^ and a high-throughput immunomagnetic microfluidic device for rare cell enrichment^22,23^, we systematically investigated genes promoting CTCs and tumor progression (Fig. 1a and Extended Data Fig. 1a). The *in vivo* genome-wide loss-of-function CRISPR-Cas9 screen was conducted using the prostate cancer cell line PC-3M as a model of tumor progression. After subcutaneous injection of the TKOv3-edited cancer cells (n = 3.0 ×10^7^ cells injected per animal) into the immunodeficient mice (n = 6 mice), tumor growth from the parental PC-3M cells or TKOv3-transduced PC-3M cells was monitored and compared for 3 weeks. No significant differences in the size of the primary tumors between these two groups were observed (Fig. 1b). At a predefined end point of 3 weeks, whole blood was collected from each animal to isolate CTCs for sgRNA enrichment analysis. Efficient isolation of CTCs from whole blood was accomplished by the immunomagnetic labeling of the epithelial marker EpCAM on the cell surface and magnetic deflection of CTCs away from blood cells using metglas tracks embedded in a microfluidic chip (Extended Data Fig. 1a, b) ^23^. CTCs collected were lysed ^24^ to obtain genomic DNA for the amplification of sgRNA regions using PCR and the determination of each sgRNA abundance using NGS analysis (Extended Data Fig. 1c). CTC sgRNA counts were successfully obtained for all the 6 mice tested using our *in vivo* CTC CRISPR screening pipeline, with consistent total sgRNAs counts across replicates as detailed below.

**Figure 1:**
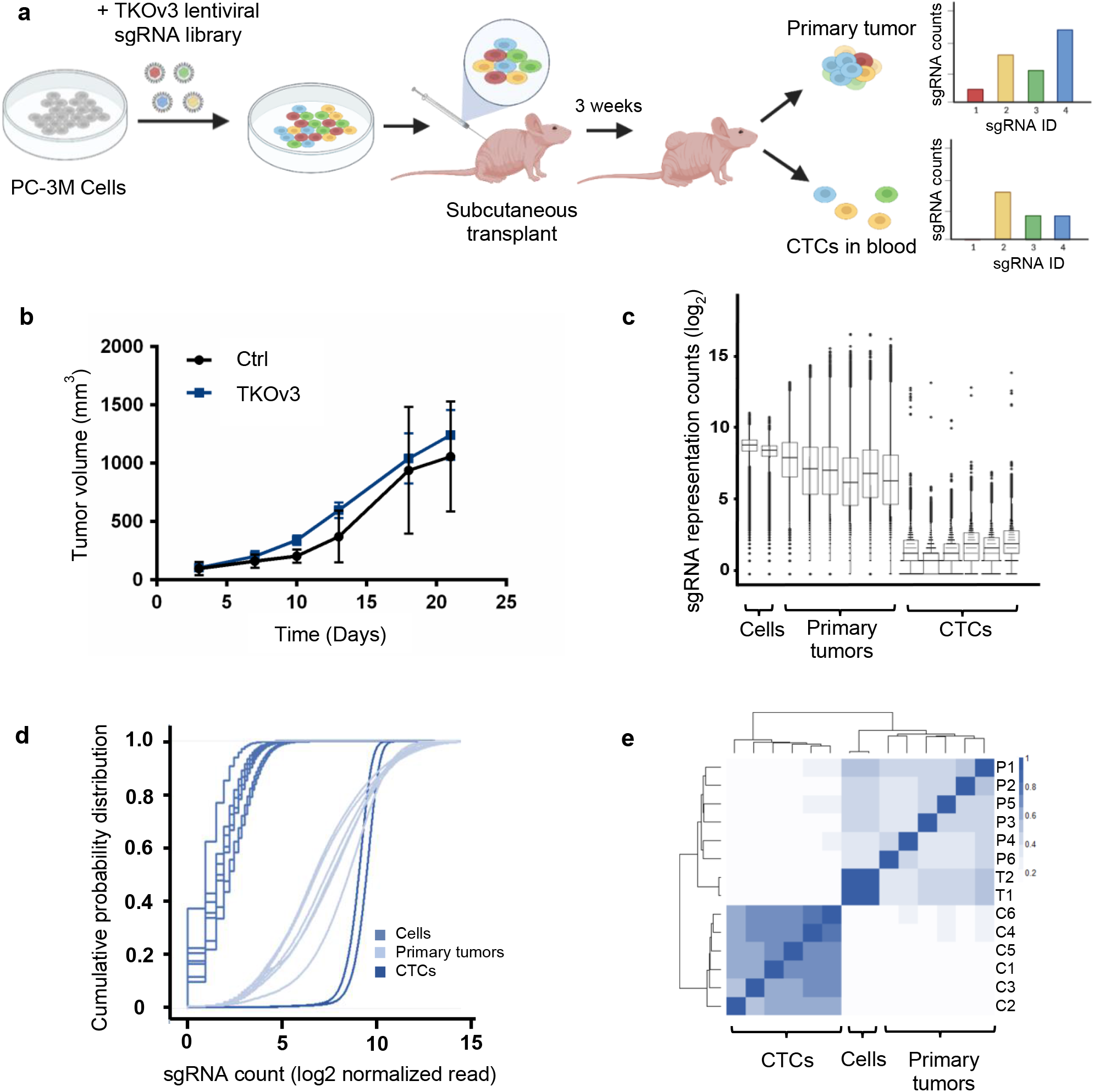
An *in vivo* genome-wide CRISPR-Cas9 knockout screen identifying CTC-promoting genetic factors. **a**. Experimental design of the genome-wide CRISPR-Cas9 KO screen focused on CTCs. **b**. Primary tumour growth curves of the immunocompromised mice subcutaneously transplanted with either the TKOv3-transduced PC-3M cells, or the control cells (n = 3 for each group). Error bars indicate SD. **c**. Box plot of the sgRNA counts from the TKOv3-transduced cell pool before transplantation (Cells), primary tumours, and CTCs. **d**. Cumulative probability distribution of sgRNA counts from the TKOv3-transduced cell pool before transplantation (Cells), primary tumours, and CTCs. **e**. Pearson correlation coefficient of the sgRNA counts from the TKOv3-transduced cell pool before transplantation (Tn), primary tumours (Pn), and CTCs (Cn).

For sgRNA enrichment analysis, we first analyzed the TKOv3 library attrition across the initial CRISPR-edited PC-3M cells, the primary tumors, and the isolated CTCs. Compared to the sgRNA counts in the initial TKOv3-transduced PC-3M cell pool before injection, majority of the sgRNAs from the library had significantly reduced or no representation in the isolated CTC samples (Fig. 1c, d). Given that 30 million cells (each contained one sgRNA) were xenografted and a library size of 71,090 sgRNAs, an initial 400x representation of the TKOv3 library was obtained. With a median capture of 100 CTCs per animal, a drastic library attrition at the CTC level was expected. Notably, the reduction of the sgRNA representation was much less in the primary tumor samples since substantially more cells and input materials were available for NGS (Fig. 1c, d and Extended Data Fig. 2a). In addition, the overall distribution of sgRNAs in the CTC samples clustered tightly with each other and formed a clade distinct from the sgRNAs representing the CRISPR-edited PC-3M cells and primary tumor samples (Fig. 1e). When individual sgRNA enrichment was examined for the primary tumors, a clear trend emerged with the top-ranked genes representing well-established tumor suppressor genes (TSGs) such as *TSC1* ^25,26^ (Supplementary Table 1). Frequent loss-of-function mutations of the top gene hits were also observed in patient tumor data collected from COSMIC with copy number variation analysis^27,28^ (Extended Data Fig. 3).

### Selection of top enriched sgRNAs in CTCs

In order to analyze the CTC-enriched sgRNAs, we developed two selection criteria for robust candidate hit identification (Fig. 2a, 2b, 2c). In the first round of selection, we prioritized sgRNAs that were consistently enriched in CTC samples harvested across all the 6 mice injected. This resulted in the identification of 149 sgRNAs that were enriched across all the CTC samples compared to their corresponding primary tumors (Fig. 2a). The second selection criterion took into account the expected number of CTCs captured at the time of sample harvesting (median CTC count 100 per mouse). As a result, only the top 100 sgRNA counts from each of the 6 replicates were pooled together to generate a list of 162 unique sgRNAs, excluding the repetative sgRNAs (Fig. 2c). After filtering for sgRNAs that satisfied both selection criteria, the final CTC sgRNA enrichment list consisted of 38 unique sgRNAs and the genes targeted by these sgRNAs were selected for downstream analysis (Fig. 2b, d). The biological pathways represented by these 38 genes were found to be highly related to metastatic processes such as the regulation of apoptosis and cell proliferation (Fig. 2e). Since no more than 1 sgRNA was enriched for each of the 38 genes selected from the genome-wide screen, we designed a sub-pool CRISPR KO library to functionally validate the top hits from the initial genome-wide screen and a true top hit must have multiple sgRNAs present in the top enriched sgRNA list generated from the sub-pool screen.

**Figure 2:**
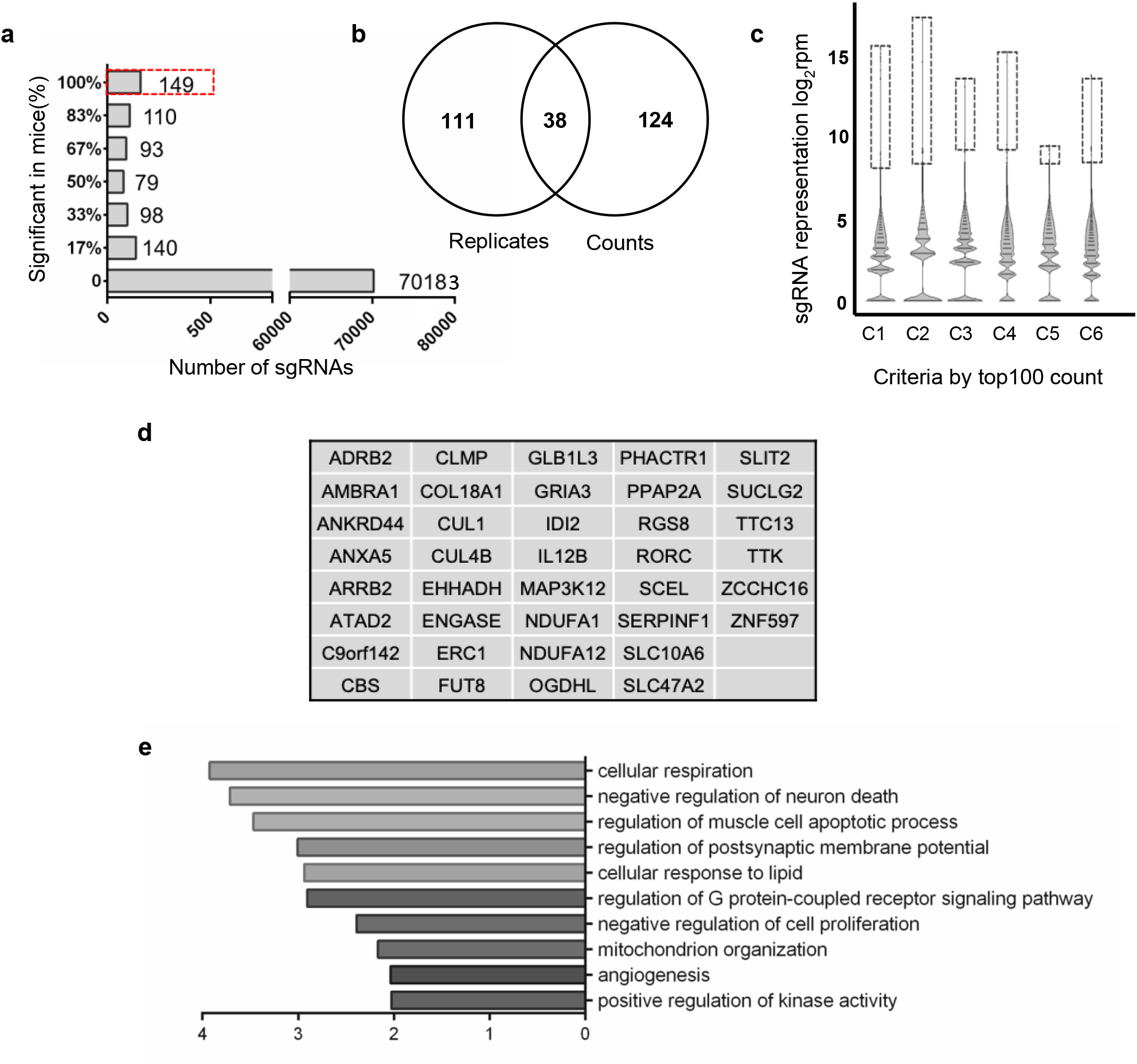
Identification of top hits from the genome-wide CRISPR-Cas9 KO screen. **a**. Frequency (percent of mice) of each sgRNA being enriched in CTCs compared to the corresponding primary tumours. The dashed box highlights the 149 sgRNAs enriched in CTCs across all 6 mice. **b**. Venn diagram showing selection of the top-enriched sgRNAs from the CTC samples. 38 sgRNAs satisfied both the “counts” criteria in (**c**) and the “replicates” criteria in (**a**), and their targeted genes were selected for downstream analysis. **c**. Violin plots of the sgRNA counts from the CTC samples. The dashed boxes highlight the top 100 sgRNA counts from each CTC sample. **d**. List of the 38 genes selected for the construction of the sub-library. **e**. Biological pathways being enriched from the 38 genes selected.

### Functional validation of top hits with sub-library screen

A customized sub-pool CRISPR KO library consisting of 400 sgRNAs specifically targeting the 38 genes of interest (10 sgRNAs per gene) plus 20 non-targeted control sgRNAs was generated. CRISPR-edited PC-3M cells using the sub-pool library were xenografted using an analogous protocol to the original screen, including the number of mice used (n = 6 mice) and the number of cells injected (n = 3.0 ×10^7^ cells injected per animal) (Extended Data Fig. 1d). CTC capture was improved by viral transducing the GPI-anchored Myc-tag into PC-3M cells before the sub-pool screen, allowing Myc-based immunomagnetic capture of CTCs (Extended Data Fig. 1b). As observed previously, compared to the other sample groups (plasmid pool, infected cells before transplant, primary tumors), the number of unique sgRNAs with significant representation in the total sgRNA reads was dramatically reduced for the CTCs (Fig. 3a and Extended Data Fig. 2b). Notably, reads for the sgRNAs targeting *SLIT2* accounted for 10% of the total sgRNA counts in the CTC samples, with 4 out of the 10 sgRNAs targeting *SLIT2* highly represented in the CTC population (Fig. 3b). Accordingly, *SLIT2* KO was ranked to be the most significant loss-of-function mutation that appeared to increase the number of CTCs, with a much higher ranking-score than the rest of the genes tested in the sub-library (Fig. 3c). *SLIT2* was thus selected for downstream analysis.

**Figure 3:**
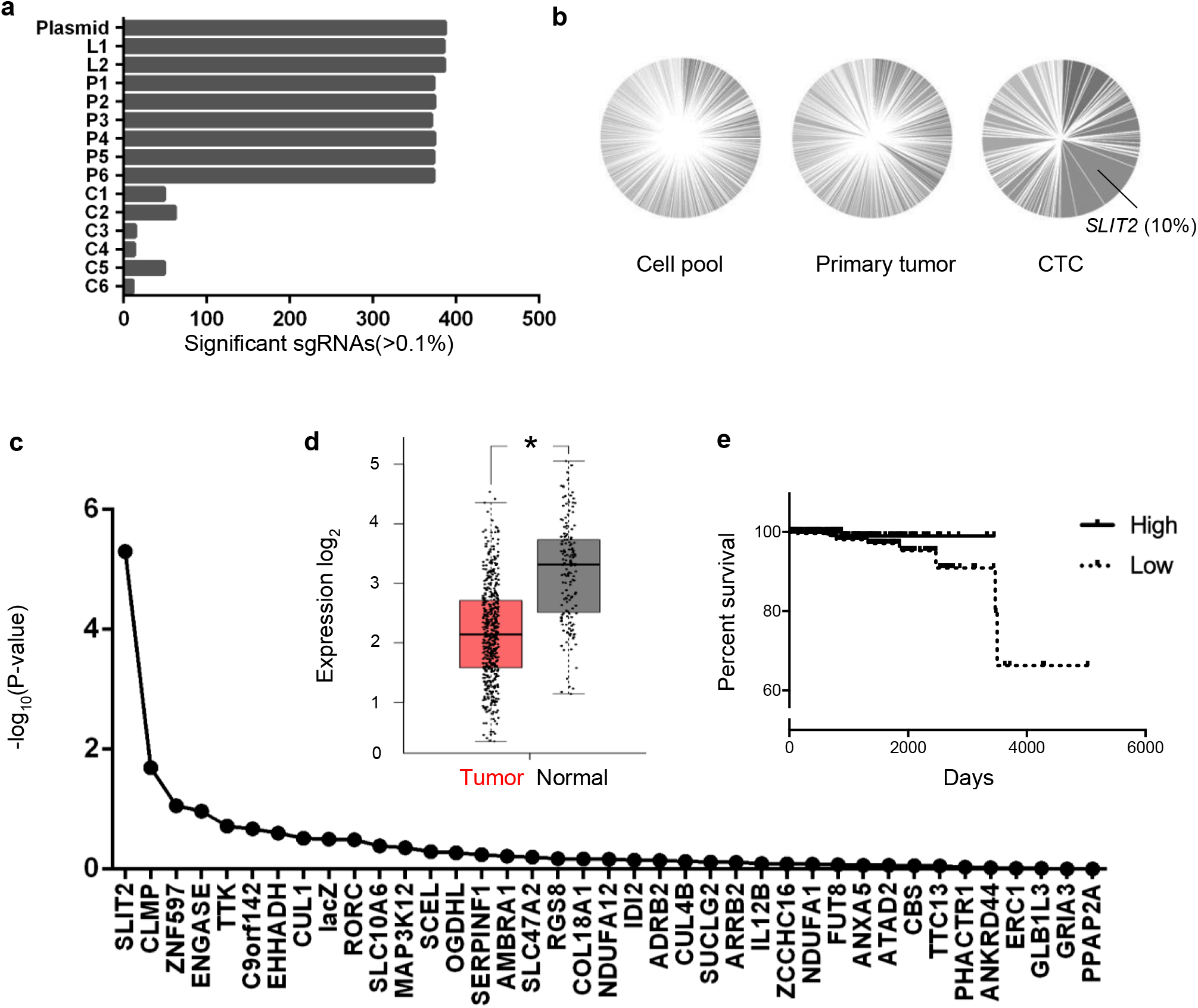
Loss of *SLIT2* was a key factor of CTC production in the sub-pool screen and was associated with poor prognosis of prostate cancer patients. **a**. Bar graph of the number of significant sgRNAs (counts > 0.1% of total reads) from the plasmid library (Plasmid), the initial cell pool before transplant (Ln), primary tumors (Pn), and CTCs (Cn) of the sub-library CRISPR KO screen. **b**. Pie chart showing gene enrichment (percentage of total reads) of the initial cell pool, primary tumor, and CTC samples from the sub-pool CRISPR KO screen. **c**. Gene ranking from the sub-pool CRISPR KO screen. *SLIT2* KO was found to be the most abundant gene KO present in CTCs. **d**. *SLIT2* expression levels in tumor and normal tissues of prostate adenocarcinoma (PRAD) patients were compared using RNAseq data collected from TCGA and prepared by GEPIA2. (Tumor: n = 492; Normal: n= 152) p-value cutoff was set to be 0.01 *P<0.01 **e**. Survival plot of PRAD patients with low or high expression of *SLIT2*. Data was collected from TCGA, with an expression cut-off at 2.45 FPKM. (High expression: n = 189; Low expression: n= 305)

### Loss of *SLIT2* is associated with poor prognosis of prostate cancer patients and induced metastasis *in vivo*

SLIT2 belongs to a family of secreted proteins with a known role in axon guidance through interaction with the Roundabout (ROBO) family of receptors ^29,30^. *SLIT2* was also found to be expressed in various cancer tissues with context-dependent promotion or inhibition of cancer progression ^31–33^. However, no previous studies had focused on the expression of *SLIT2* in CTCs and the effect of this genetic factor on generation of CTCs, especially in a context of prostate cancer. In the light of our CTC CRISPR KO screen conducted with the prostate cancer cell line PC-3M, we hypothesized that *SLIT2* negatively regulates CTC production and thereby prevents metastatsis.

Based on patient data collected from The Cancer Genome Atlas (TCGA), *SLIT2* expression level is significantly reduced in tumor tissues compared to normal prostate tissues (Fig. 3d). Low *SLIT2* expression is also associated with poor prognosis and overall survival of prostate cancer patients (Fig. 3e). Based on COSMIC, of the 1068 patients with information regarding the functional status of *SLIT2*, 65% of patients possessed *SLIT2* loss-of-function mutations. In addition, reduced expression of *SLIT2* in tumor tissues compared to the corresponding normal tissues is observed in 26 out of the 31 human cancers recorded in TCGA (Extended Data Fig. 4), indicating the generality of the negative correlation between *SLIT2* expression and development of tumors.

To investigate the role of *SLIT2* in a prostate cancer model, *SLIT2* KO PC-3M cells were generated using CRISPR editing to probe the effects of losing this gene *in vivo* (Extended Data Fig. 5a). Equal numbers of non-targeted control (NTC) or the *SLIT2* KO PC-3M cells (1 × 10^6^) were orthotopically injected into the prostate of immunodeficient mice. At the end point of tumor growth, the number of CTCs present in the whole blood of the mice injected with *SLIT2* KO cells was increased up to fivefold compared to the group injected with the NTC cells (Fig. 4a and Extended Data Fig. 5d, e). The xenograft generated from *SLIT2* KO cells also exhibited enhanced proliferation of cancer cells, resulting in significantly increased tumor size compared to the control group (Extended Data Fig. 5b, c). In order to assess if the enhanced CTC production in *SLIT2* KO model was solely because of elevated tumor size, *SLIT2* KO xenografts with comparable tumor sizes to the control were generated with a decreased number of *SLIT2* KO cells at injection (Fig. 4b). In this case, *SLIT2* KO again exhibited a significantly increased number of CTCs in whole blood compared to the control group (Fig. 4b), indicating enhanced intravasation and dissemination of cells lacking *SLIT2*. Since less cells were used for injection and blood samples were harvested earlier in this case to ensure comparable tumor sizes between control and *SLIT2* KO cells, overall CTC counts were lower compared to the previous injection. In addition, severe infiltration of neoplastic cells was observed inside the lymph nodes of the *SLIT2* KO models, with cases of subcapsular and intravascular invasion found in multiple H&E-stained histology sections (Fig. 4c, d). No invasion or peripheral infiltration of neoplastic cells was observed for the prostate cancer model generated from the control cells at the time of harvest. Lymph node was used as a representation of metastasis model based on our previous observation that PC-3M had more chances to develop metastasis at the lymph nodes at late metastatic stage compared to other prostate cancer cell lines such as LNCaP ^34^. We concluded that knockout of *SLIT2* promotes CTC numbers and thereby increases metastatic burden in a prostate cancer model.

**Figure 4:**
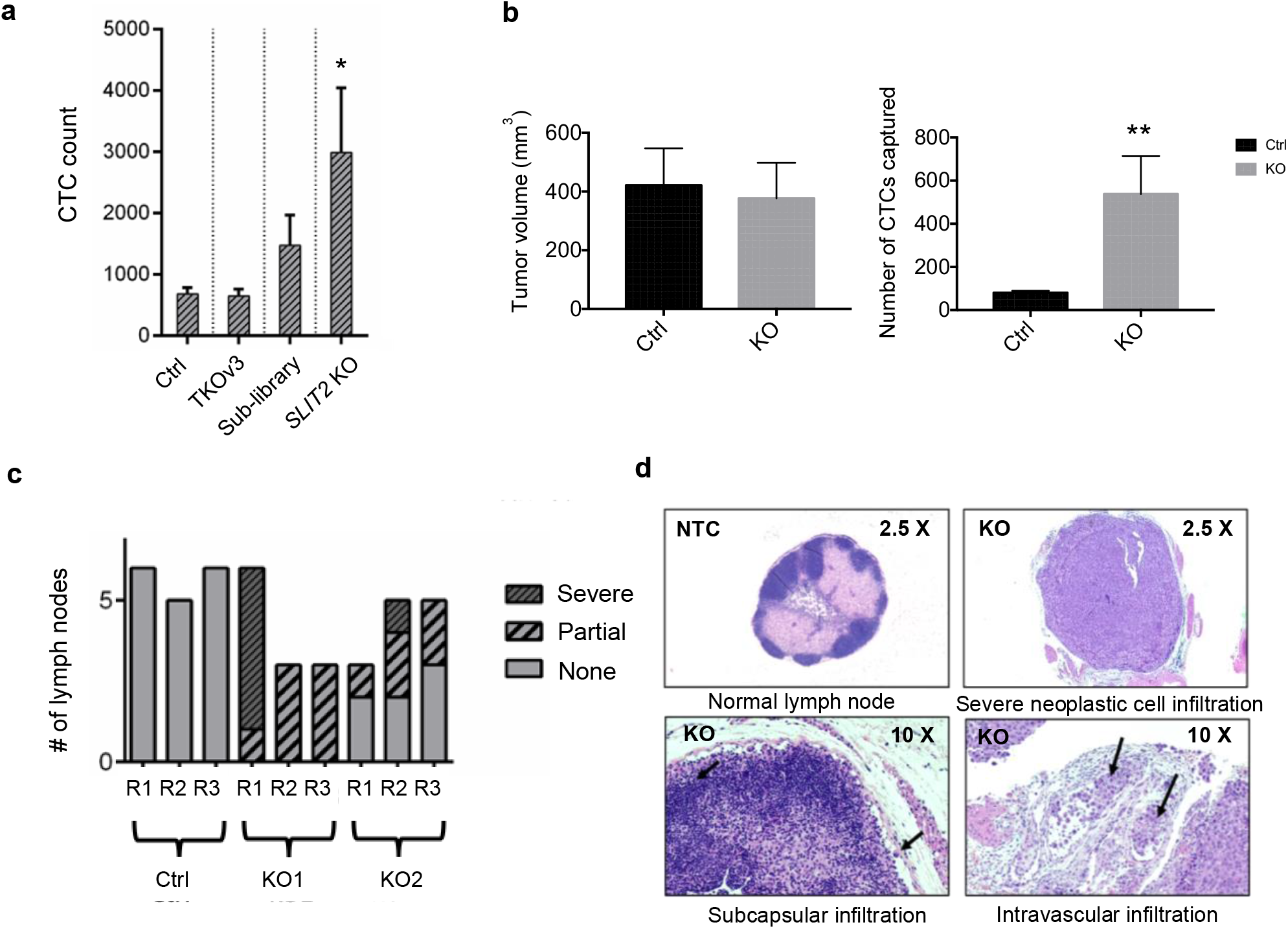
SLIT2 KO promoted cancer progression *in vivo*. **a**. Bar graph of the number of CTCs counted from the whole blood of the immunocompromised mice after transplant of non-targeted control, genome-wide CRISPR KO library (TKOv3), tumor suppressor gene-enriched CRISPR KO sub-library, or *SLIT2* KO cells (n = 3 for each group). Error bars indicate SD. P value was calculated by two-tailed unpaired t-test. P=0.045. **b**. Bar graph comparing CTC number resulted from similar tumor size of NTC and *SLIT2* KO cells (n=3 for each group). Error bars indicate SD. P value was calculated by two-tailed unpaired t-test. P=0.0072. **c**. Bar graph of the number of lymph nodes with normal, neoplastic cell infiltrated, and severely infiltrated tissues. Histology slides were prepared from lymph nodes collected from the immunocompromised mice after orthotopic injection of either non-targeted control or the two *SLIT2* KO cell lines. **d**. Representative images of the H&E stained sections showing normal lymph node tissues (NTC), as well as severe, subcapsular, and intravascular neoplastic cell infiltration (*SLIT2* KO).

### Loss of *SLIT2* results in enhanced migratory and invasive behavior of prostate cancer cells

To understand mechanisms by which *SLIT2* modulates the metastatic process, we studied *SLIT2* KO and NTC cells using a variety of phenotypic assays to assess cell proliferation, deformability, and invasion. 3D spheroids cultured from the *SLIT2* KO and the NTC cells were comparable in size and composed of similar number of cells, but the *SLIT2* KO spheroids appeared to be poorly organized with cells on the outer surface of the spheroids being loosely attached (Fig. 5a and Extended Data Fig. 6). To evaluate the impact of *SLIT2* loss-of-function on the migratory potential of PC-3M cells, a specialized microfluidic device with migration channels connecting the cell loading and the nutrient holding pools was designed (Extended Data Fig. 7). The channels had a constant height of 5 μm, but decreasing width thereby requiring the cells to undergo remodelling to squeeze through narrowing channels towards the nutrients (Extended Data Fig. 7). While a comparable number of *SLIT2* KO and NTC cells traveled through the migration channels wider than 10 μm, significantly more *SLIT2* KO cells were able to fit into the channels of width 10 μm or less (Fig. 5b, Extended Data Fig. 7b).

**Figure 5:**
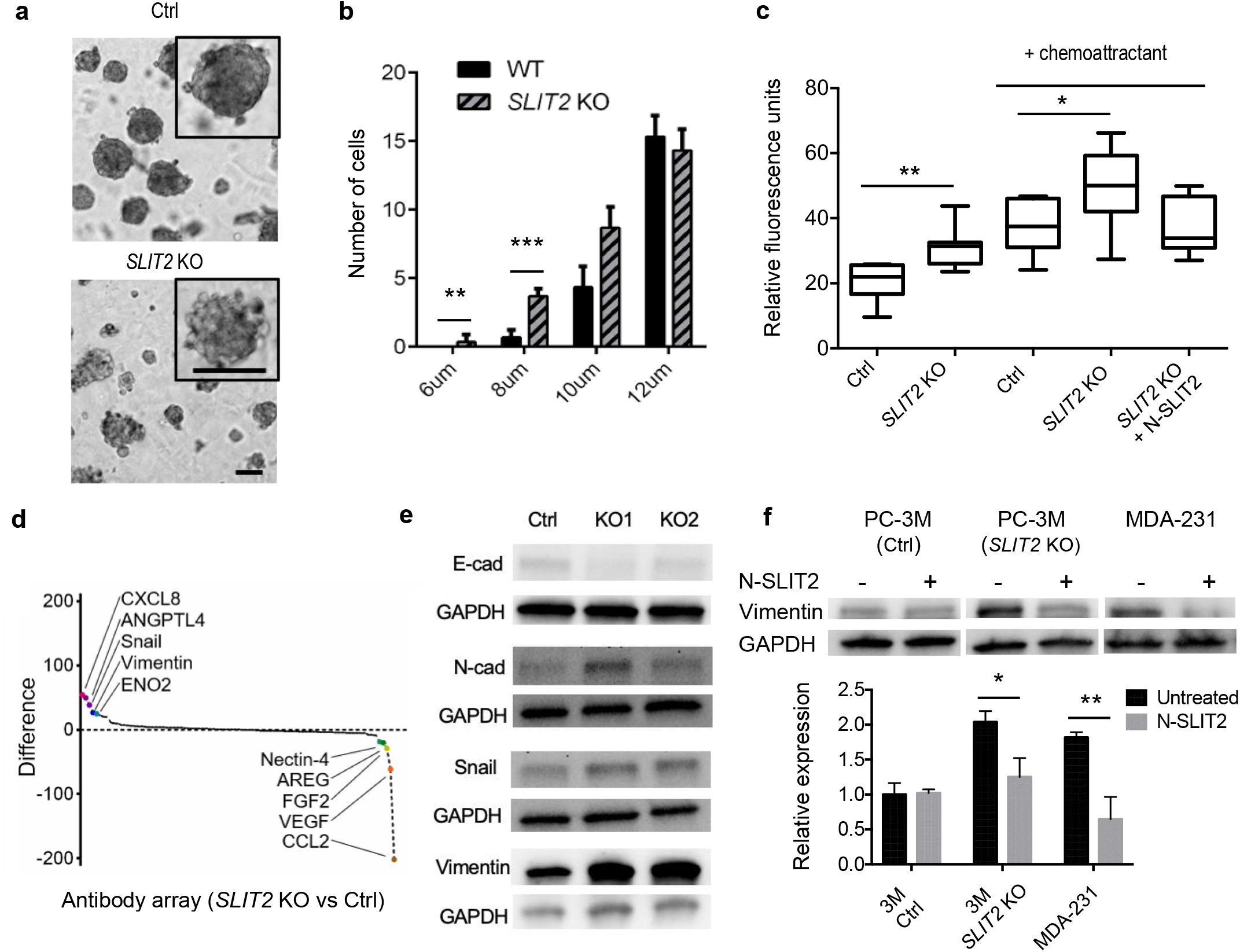
*SLIT2* KO facilitated metastasis with activation of EMT. **a**. Representative images of the spheroids formed from non-target control (NTC) or *SLIT2* KO PC-3M cells. **b**. Bar graph comparing the number of deformed NTC versus *SLIT2* KO cells at different migration channel width (n = 3 for each group). **c**. Box plot comparing the number of NTC versus *SLIT2* KO cells invaded across the transwell membrane with or without the presence of the chemoattractant or N-SLIT2 protein in the culture media (n = 8 for each group). **d**. Proteins ranked based on levels of differential expression in *SLIT2* KO compared to NTC cells as detected by the oncology antibody array. **e**. Immunoblots detecting E-cadherin, N-cadherin, Snail, and Vimentin expression in *SLIT2* KO and NTC cells. **f**. Immunoblots detecting Vimentin protein levels in control and *SLIT2* KO PC-3M, as well as MDA-231 cells, either with or without supplementation of N-SLIT2 protein in culture medium. Error bars indicate SD. P values were calculated by two-way ANOVA, except for (**c**) which was by two-tailed unpaired t-test. ***P<0.001; **P<0.01; *P<0.05; NS:P>0.05.

The invasive behaviour of *SLIT2* KO and control cells was also tested using a transwell cell invasion assay. A larger number of *SLIT2* KO cells migrated from the top to the bottom of the membrane compared to the control group, with or without the presence of the chemoattractant in the lower chamber (Fig. 5c). Importantly, the number of invading *SLIT2* KO cells was similar to control cells when the media was supplemented with exogenous SLIT2 protein (Fig. 5c). Importantly, reuslts of cell deformability and invasion assays were independent of cell proliferation rates, as no significant difference between the cell numbers of control and *SLIT2* KO was found after 2D culturing for 3 days (Extended Data Fig. 8).

### Loss of *SLIT2* results in activation of epithelial-to-mesenchymal transition

The interrogation of a panel of cancer-related proteins using an oncology antibody array revealed several strong hits that connected the migratory and invasive phenotypes observed in *SLIT2* KO cells to the epithelial-to-mesenchymal transition (EMT) (Fig. 5d and Extended Data Fig. 9). Considering the most elevated protein family or biological pathways identified from the array, protein levels of both mesenchymal markers Vimentin and Snail were enhanced in *SLIT2* KO cells. Thus, we hypothesized that the knockout produced a more mesenchymal phenotype. Reduced expression of the epithelial marker (E-cadherin) and increased expression of mesenchymal markers (Vimentin, N-cadherin, Snail) were observed for *SLIT2* KO cells compared to NTC cells, indicating that EMT may be linked to the metastatic phenotype of cells lacking *SLIT2* (Fig. 5e). Expression level of the epithelial marker EpCAM on the cell surface was also checked as it was used for the CTC capture during the screening process and no significant difference was found comparing control and *SLIT2* KO cells (Extended Data Fig. 10).

Reversal of the mesenchymal phenotype was explored in a rescue experiment where exogenous SLIT2 was added to culture media and the level of Vimentin was analyzed. Addition of SLIT2 did not have a significant effect on Vimentin expression in control PC-3M cells (Fig. 5f). However, the level of Vimentin in *SLIT2* KO PC-3M cells, which was originally doubled compared to the control cells, was significantly reduced with the addition of exogenous SLIT2 (Fig. 5f). In addition, we also tested Vimentin levels for a metastatic breast cancer cell line (MDA-MB-231), and significant decrease of Vimentin expression was observed with the presence of SLIT2 (Fig. 5f), extending the link between *SLIT2* and EMT in a different context.

### Loss of *SLIT2* resulted in elevated complex I expression with hypersensitivity to rotenone

To elucidate additional drivers of the invasive phenotype of *SLIT2* KO cells, gene expression profiling of NTC and *SLIT2* KO cells was also performed (Fig. 6a, b). Strikingly, *SLIT2* KO cells exhibited increased expression of one component of the ATP synthase (*ATP6*) and multiple subunits of the complex I (*MT-ND1*, *MT-ND2*, *MT-ND5*) belonging to the mitochondrial electron transport chain (ETC), making metabolic process and oxidative phosphorylation the most enriched gene ontology term (Fig. 6b and Extended Data Fig. 11). Elevated expression of ETC components suggested a possible dependency of *SLIT2* KO cells on mitochondrial activity and oxidative metabolism. Confirming this prediction, *SLIT2* KO cells were significantly more sensitive to treatment with the complex I inhibitor rotenone when compared to the control cells (Fig. 6c). To test if the elevated activity of complex I contributed to the increased migratory potential of the PC-3M cells lacking *SLIT2*, we tested the ability of rotenone to block migration of *SLIT2* KO and control cells using the microfluidic device described above (Extended Data Fig. 7). Upon treatment of rotenone, significant reduction of the number of deformed cells was observed in both groups at wider channel width (Fig. 6d). However, for the narrower channels, rotenone was more effective in inhibiting the migration of the *SLIT2* KO cells than on the NTC cells (Fig. 6d). Importantly, other two prostate cancer cell lines LNCaP and PC3 also showed enhanced sensitivity to rotenone with *SLIT2* being knocked out (Extended Data Fig. 12 and 13). Treatment of rotenone significantly reduced the invasiveness of LNCaP and PC3 with *SLIT2* KO (Extended Data Fig. 14).

**Figure 6:**
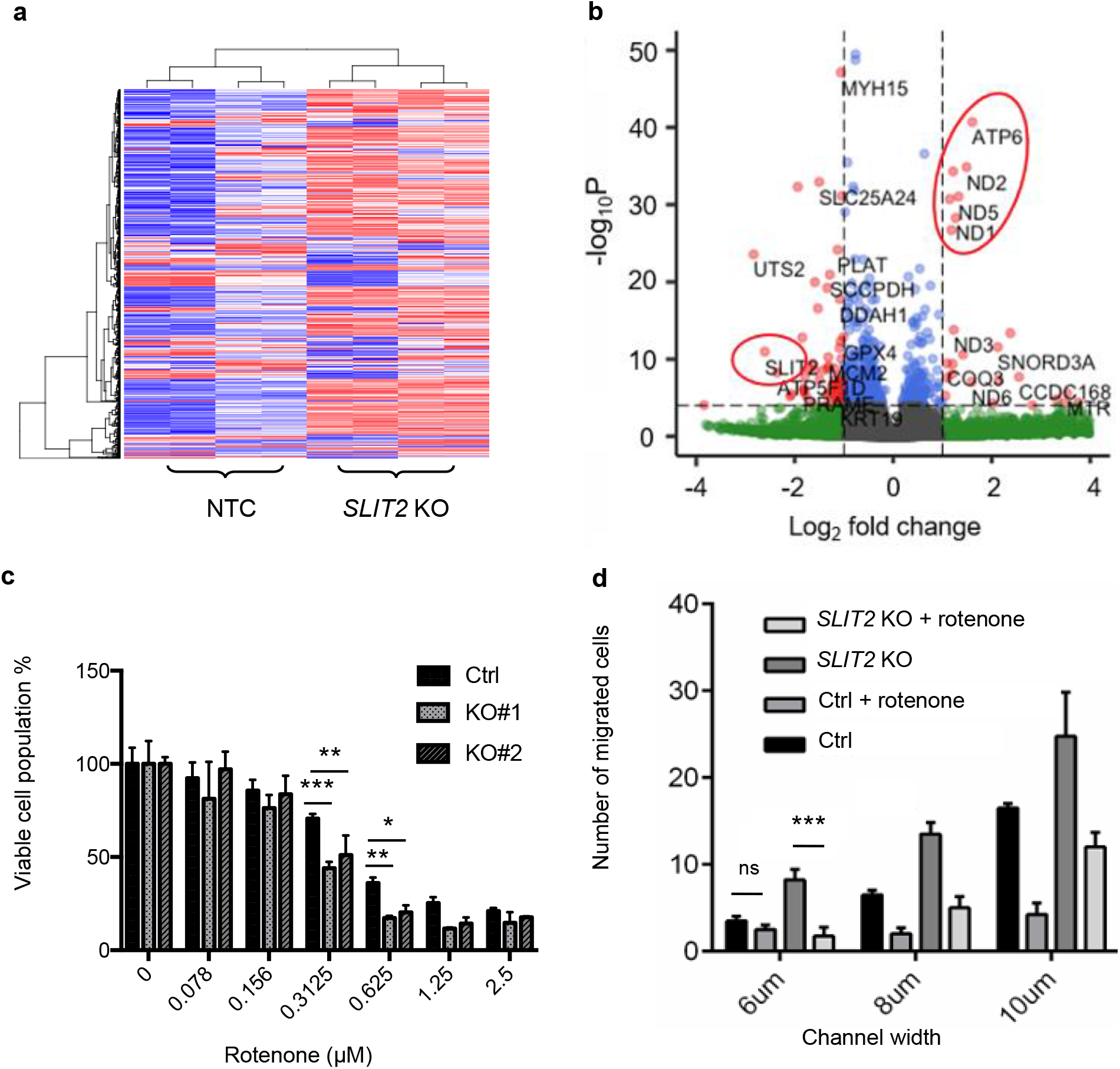
*SLIT2* KO resulted in elevated complex I expression and hypersensitivity to rotenone. **a**. Heat map of the top 1000 differentially expressed genes in *SLIT2* KO cells compared to the non-targeted control. **b**. Volcano plot highlighting (in red circles) the most differentially expressed genes of *SLIT2* KO cells compared to NTC. **c**. Bar graph showing viability of the *SLIT2* KO compared to NTC cells under titration of rotenone (n = 3 for each group). **d**. Bar graph showing the number of *SLIT2* KO cells migrated across various channel width compared to the NTC cells, with or without the presence of rotenone at IC20 (n = 3 for each group). Error bars indicate SD. P values were calculated by two-way ANOVA. ***P<0.001; **P<0.01; *P<0.05; NS:P>0.05.

## Discussion

Previous studies of the role of *SLIT2* in metastasis has produced contradictory observations in different experimental systems, with limited studies on prostate cancer models ^32,33,35–38^. Promoter hypermethylation was previously found to be responsible for the repression of *SLIT2* and was associated with aggressive prostate, breast, and lung cancers ^32,33^. Importantly, a recent study showed enhanced expression of *SLIT2* in endothelial cells but repressed expression due to hypermethylation in the tumor compartment, suggesting the differential expression profiles of *SLIT2* at the tumor microenvironment as a driver for the release of tumor cells into the blood vessels ^32^. As a top hit in our genome-wide CTC CRISPR KO screen as well as in a sub-library screen, *SLIT2* was identified as a strong modulator of invasiveness and metastasis in prostate cancer. The metastasis-promoting effect of *SLIT2* KO was also confirmed in a prostate cancer mouse model in our study. No pathways have previously been connected with the invasive behaviour of prostate tumor cells carrying a loss-of-function mutation in *SLIT2*, except that the expression of *SLIT2* is regulated by the transcriptional repressor EZH2 ^33^. Our EMT and transcriptome studies revealed aberrant expression of multiple EMT markers and ETC components in *SLIT2* KO cells that potentially contribute to metastasis. Mitochondrial activity has been linked to cancer progression in various studies ^39–43^, with inhibitors of complex I reported to be promising anti-neoplastic therapies ^44,45^. Our studies suggest that cancer cells lacking *SLIT2* expression exhibit increased sensitivity towards complex I inhibition, revealing a potential novel therapeutic strategy for cancer patients with *SLIT2* loss-of-function mutations.

Bulk sequencing of tumors has uncovered a number of cancer-causing mutations, some of which were harnessed for the development of new targeted therapeutic agents. The identification of gene mutations that are functionally implicated in tumor progression is paramount to nominate new targets for cancer. One important limitation is assessing the function of genes in rare cell populations, such as in CTCs or subsets of immune cells present in the tumor microenvironment. In this study, we performed for the first time an *in vivo* genome-wide CRISPR-Cas9 KO screen targeting rare CTC cells with prostate cancer origin. Development of this experimental pipeline is transferrable and can be easily applied to search for the unique CTC-promoting factors in other cancer types. This platform can also be further applied to explore genetic factors affecting other rare cells such as immune cells or other cell types composing the tumor microenvironment. Building on this study, we anticipate that combining efficient capture of rare cells with gene editing methodologies will provide new insights into the progression of various disease, including cancer, and will lead to the imagination of new therapies.

## Methods

### Cell Culture

Prostate cancer cell line PC-3M was obtained from Dr. Alison Allan at London Health Sciences, London, ON, Canada, and was cultured in RPMI-1640 (Wisent) supplemented with 10% fetal bovine serum (Wisent) and 1% penicillin/streptomycin (Wisent), at 37 °C, 5% CO_2_. MDA-MB-231 (ATCC) and lenti-X 293T cells (Takara Bio) were grown in DMEM (Sigma) supplemented with 10% fetal bovine serum (Wisent), 4 mM stable glutamine (Sigma), and 1% penicillin/streptomycin (Wisent), at 37 °C, 5% CO_2_.

### Lentiviral production

All lentivirus was produced using lenti-X 293T cells. For TKOv3 genome-wide library production, 8.0 μg TKOv3 pooled plasmid (Addgene) was co-transfected into 80% confluent lenti-X 293T cells with 4.8 μg psPAX2 (Addgene) and 3.2 μg pMD2.G (Addgene) per 15 cm plate (Sarstedt). The mixture of plasmids (16.0 μg in 50 μL ddH_2_O) was added to 48 μL of X-tremegene 9 DNA transfection reagents (Roche) that pre-incubated for 5 min in 750 μL OptiMEM (Gibco). The mixture of plasmid-transfection reagent-OptiMEM was further incubated for 25 min before adding dropwise to lenti-X cells. 18 hr later, media was changed to fresh DMEM. Viral particles were harvested 48 hr after the media exchange, snap freezed in liquid nitrogen, and stocked at −80 °C. The customized sub-library (plasmid pool synthesized at GenScript) lentivirus was prepared using the same protocol as the TKOv3 library. Individual *SLIT2* CRISPR KO lentivirus was generated following the polyethyenimine (PEI) transfection protocol (Addgene). sgRNAs targeting the *SLIT2* gene were cloned into the lentiCRISPRv2 (Addgene) and the 20 bp sgRNA sequences used were CGGGTTGGTCTGTACACTCA for KO 1 and ACGGAAAGCTTTCCTTGGGA for KO 2. For the non-targeted control, the sgRNA sequence used was CCCGAATCTCTATCGTGCGG.

### Lentiviral transduction and MOI determination

Lentivirus was titrated into 2 × 10^5^ PC-3M cells on 6-well plates with 2 mL complete RPMI-1640 supplemented with 8 μg/mL polybrene (Sigma). After overnight infection, media was exchanged for fresh RPMI supplemented with 2 μg/mL puromycin (Sigma). After 48 hours of antibiotic selection, multiplicity of infection (MOI) was determined by comparing the number of cells remained at specific viral load with or without puromycin selection using 0.1% crystal violet staining (Sigma) and absorbance reading at 590 nm. A MOI of 0.3 was used for the transduction of CRISPR libraries into the PC-3M cells.

### Genome-wide CTC CRISPR KO screen

1.0×10^8^ PC-3M cells were transduced with TKOv3 library (71,090 sgRNAs) at a MOI of 0.3 using polybrene as transduction reagent as described in the previous section. After 2 rounds of puromycin selection, 3 × 10^7^ transduced cells were subcutaneously injected into the flank of each of the 6 male athymic nude (nu/nu) mice of 6-8 weeks old (1.5 × 10^7^ each side per mouse). Tumor volume was measured twice a week using a caliper and calculated using the formula length x width^2^/2. Mice were euthanized 3 weeks after transplant to collect tumor and whole blood samples (cardiac puncture). All animal work was conducted following the protocol approved by University of Toronto Animal Care Committee under the guidelines of Canadian Council on Animal Care (CCAC).

### PRISM chip design and fabrication

An initial iteration of the PRISM chip ^23^ was modified to increase throughput to 3×10^7^ cells/hr/chip and sort cells into zero/low-, medium- and high-expression outlets. Magnetically labeled cells are deflected laterally based on their level of expression. Deflection is facilitated using guides made from highly magnetically permeable alloy set at an angle from the fluid flow (5° for medium and 20° for high). In the presence of a magnetic field from a permanent magnet, localized regions of high magnetic flux are found at guide edges causing the attraction of magnetic nanoparticles ^46^. A balancing of the magnetic force and the Stokes drag force from fluid flow causes magnetically labeled cells to follow the deflection guides until the angle changes or the guides end at the lateral edges of the device. Angle changes of the guides causes an increase in the component of the drag force and medium-expression cells to detach. Chips were fabricated on a soda lime glass wafer (UniversityWafer). Guides were patterned using photolithography and wet etching and channels were created using SU-8 photo resist (Kayaku AM). A PDMS slab (Dow Sylgard) was bonded to encapsulate the device according to an existing protocol ^47^. Multiple chips were setup in parallel to simultaneously isolate CTCs from multiple blood samples. To prepare the PRISM chips for CTC isolation, the chips were primed with sterile water containing 0.1 % Pluronic F68 (Sigma) under constant hydraulic pressure for overnight before use. The non-ionic surfactant Pluronic F68 prevented nonspecific capture of cells in the microchannel and the constant hydraulic pressure removed small air bubbles. For immunomagnetic-based cell sorting, the Prism chip was first mounted on a permanent magnet (BY088-N52, K&J Magnetics). Before sample loading, a buffer solution made of HBSS (GIBCO) supplemented with 2% BSA (Sigma) and 5 mM EDTA (BioShop) was introduced to the Prism chip to remove the pluronic water.

### CTC collection and cell lysis

To isolate the CTCs, each blood sample was diluted in a 1 to 1 ratio with the HBSS buffer solution and incubated with 10% EpCAM MicroBeads (Miltenyi Biotec) with shaking at room temperature for an hour. Afterwards, equal volumes of buffer solution and the blood samples containing the magnetically labeled CTCs were loaded onto the buffer inlet and sample inlet respectively before withdrawn from the chip at a flow rate of 2 mL/hr. After sample processing, 300 μL of buffer solution was loaded to both the sample and buffer inlets and withdrawn at the same flow rate to wash out remaining samples in the microchannels. Solution collected from the high-expression outlet (~500 μL) of each chip was injected into a 1.5 mL centrifuge microtube (Axygen) and set on the DynaMag™-2 magnet (ThermoFisher) for an hour. After the cells settled down, supernatant was pipetted out slowly without touching the bottom or the wall of the tube, leaving minimal solution (5-10 μL) containing CTCs. Alkaline lysis method adapted from a previous publication was used to lyse CTCs ^24^. Briefly, 10 μL 200 mM KOH with 50 mM dithiothreitol was added to CTC solution and heated at 65 °C for 10 min in a thermocycler. Lysate was then neutralized by 10 μL of 900 mM Tris-HCl pH 8.3 with 300 mM KCl and 200 mM HCl.

### Genomic DNA extraction and PCR amplification of sgRNA

To extract gDNA from the CRISPR library-transduced cell pool (3-5 × 10^7^ cells) and the tumor tissues (100-200 mg), QIAamp DNA Blood Maxi Kit (Qiagen) was used following the manufacturer’s instructions. For gDNA extraction from tumor tissues, tumors were chopped into small pieces using a razor blade and incubated with 1.8 mL tissue lysis buffer ATL (Qiagen) and 0.2 mL Proteinase K (Qiagen) at 56 °C overnight before processed using the QIAamp DNA Blood Maxi Kit. gDNA extracted from the initial cell pool and the primary tumor samples were checked for 230/280 and 260/280 ratios before sending to the Princess Margret Genomics Center (PMGC, Toronto, ON Canada) for PCR amplification of the sgRNA region and generation of the barcoded sgRNA library. For the CTC samples, gDNA extraction was not necessary and 10 μL of the neutralized lysate from each CTC sample was directly set for PCR amplification of the sgRNA region. Each 100 μL PCR reaction was set as following:

10 μL 10 x buffer (supplied with Ex Taq)
8 μL dNTP (supplied with Ex Taq)
5 μL Amp1 primer mix (10 μM)
10 μL neutralized CTC lysate
65.5 μL water
1.5 μL Ex Taq DNA polymerase (Takara Bio)

The thermocycling parameters used were:

Step 1: 95 °C for 1 min
Step 2: 95 °C for 30 sec
Step 3: 52 °C for 30 sec
Step 4: 72 °C for 20 sec
Repeat steps 2 – 4 for 34 additional times
Step 5: 72 °C for 2 min
Step 6: 4 °C hold

PCR products were run on a 1 % agarose gel to confirm the size of the product (expected size: 235 bp) before sent to PMGC for barcoded sgRNA library generation. NGS was done using Illumina NovaSeq 6000 with 1 million reads per sample.

### CTC sub-pool CRISPR KO screen

Sub-library (400 sgRNAs) was synthesized and cloned into the same vector as the TKOv3 genome-wide library (lentiCRISPRv2) by GenScript. Cloned sub-library was deep sequenced to verify sgRNA representation in the plasmid pool. Lentivirus production, MOI determination, and *in vivo* screen were conducted in the same manner as the TKOv3 library, except that GPI-anchored Myc tag was lentivirally introduced to the PC-3M cells for displaying Myc on the cell surface before introducing the sub-library to the cells. Thus, CTCs in blood samples were labeled with 1% biotinylated anti-Myc antibody (Abcam, ab197139) for an hour and 5% anti-biotin microbeads (Miltenyi Biotech) for an hour before being isolated on the Prism chip. Lysis and sgRNA PCR amplification of CTCs in the sub-library screen used the same protocol as the genome-wide screen.

### NGS data and sgRNA enrichment analyses

The genome-wide screen data was mapped by PMGC, and the customized sub-library data was mapped following the MAGeCK pipeline ^48^. Number of reads in each sample was normalized by converting raw sgRNA counts to reads per million (rpm). The rpm values were then subject to log2 transformation for downstream analyses. To generate correlation heatmaps, NMF R package was used ^49^ and Pearson correlations between individual samples were calculated using log2 rpm counts. Empirical cumulative distribution plot was generated using the ecdfplot function in the latticeExtra R package. Biological pathway enrichment analysis was performed using Metascape ^50^. For sgRNA enrichment analysis, fold change was obtained using MAGeCK and two criteria were used to identify the top candidate genes from the genome-wide screen based on CTC sgRNAs: 1) if a CTC sgRNA was enriched compared to its counterpart in the corresponding primary tumor sample (fold change>=1.0) across all the samples (n = 6); 2) if a sgRNA was counted as the top 100 in the normalized mapping results of each CTC sample. For the sub-library data, sgRNA counts were normalized and analyzed for enrichment using DrugZ ^51^.

### *In vivo* validation of screen top hit

PC-3M NTC or *SLIT2* KO cells (1.0×10^6^ each) were orthotopically injected into the prostate via the right dorsolateral lobe of 6-8 week-old male athymic nude (nu/nu) mice. After three weeks, mice were euthanized to collect primary tumor (size comparison), blood (CTC counting), and lymph nodes (metastasis histology). Lymph nodes for histology were formalin-fixed, paraffin-embedded, sectioned, and stained with hematoxylin and eosin (prepared by Pathology Research Program Laboratory at University Healthy Network). Prepared tissue sections were imaged and examined by a pathologist at Charles River laboratory for the assessment and grading of the metastatic lesions.

### Fabrication and performance of the 8-zone CTC capture device

8-zone profiling chips were fabricated using 3D-stereolithography for the 3D printed molds and standard soft lithographic techniques for subsequent steps as described previously ^52–54^. Prior to usage, the microfluidic chips were treated with 0.1% Pluronic F-68 overnight. Devices were assembled with 2 rectangular arrays of N52 neodymium magnets above and below before connected to a syringe pump (Chemyx) set to withdraw for the duration of the sample processing. Blood samples were prepared following the same protocol as for the PRISM chips before loaded onto the 8-zone device, and CTCs were captured and profiled for EpCAM expression. 500μL of PBS was withdrawn through the chip to displace the Pluronic F-68 solution before sample loading. The samples were processed at a flow rate of 750 μL/hr. The samples were then fixed with 150μL of 4% paraformaldehyde (Sigma) in PBS, permeabilized with 150μL of 0.2% Triton-X-100 (Sigma) in PBS. The chips were washed in between each step with 125μL of CliniMACS PBS-EDTA solution (Miltenyi). Captured cells were immunostained by 200μL of an antibody cocktail containing 30 μg/mL anti-pan cytokeratin AF488 (Abcam, ab277270), 6 μg/mL anti-cytokeratin 18 FITC (LifeSpan BioSciences, LS-C46335), 30 μg/mL anti-cytokeratin 19 AF488 (BioLegend, 628508), and 3 μg/mL anti-mouse CD45 APC (BD Bioscience, 559864) in PBS supplemented with 1 % BSA and 0.1 % Tween 20 (Sigma) at a flow rate of 200 μL/h. The captured cells were then stained with DAPI (Thermo Fisher, R37606) before fluorescently imaged and scanned using Nikon Ti-E Eclipse microscope with Andor’s Neo sCMOS camera. CTCs were identified as nucleus present and EpCAM^+^ CK^+^ CD45^−^ cells.

### Spheroid assay

3D spheroids consisted of NTC or *SLIT2* KO cells were grown following a protocol described previously ^55^. Briefly, RPMI-1640 was supplemented with 0.75 % methylcellulose (Sigma), and the spheroids were cultured in pre-treated ultra-low attachment plates (Corning). To determine the amount of cells consisting the spheroids in each well, CellTiter-Glo 3D Cell Viability Assay (Promega) was used following manufacturer’s instructions. Briefly, assay reagent was mixed vigorously with cell culture medium at a 1 to 1 ratio after the plate was brought to room temperature. After 25 min of incubation at room temperature, luminescence was recorded for each well.

### Migration chip fabrication and cell migration/deformability assay

The migration chip includes a sample loading channel in the middle and two stimuli loading channels on each side. There are numerous narrow migration channels (constant height at 5 μm) with varying widths ranging from 6 μm to 20 μm connecting the sample channel and stimulation channels for cell migration. The chip consists of a PDMS substrate with microchannels and a glass cover. The PDMS substrate was fabricated using soft lithography. Briefly, the liquid base and reagent of PDMS was fully mixed at a ratio of 10:1 and casted on a SU8 negative photoresist-patterned silicon mold, and then incubated at 70 °C for 2 hours for polymerization. The solidified PDMS was peeled off the mold and bonded with the glass cover upon plasma treatment. For cell deformability assay, the migration chip was first filled with 70% ethanol and UV-treated. The degassed and sterilized chip was then washed with PBS followed by cell culture medium. 3×10^6^ PC-3M NTC or *SLIT2* KO cells in 500 μL complete medium were loaded to the sample channel and incubated at 37 °C for overnight to allow cell attachment. The next day, the sample channel was washed with PBS and loaded with serum-free medium. The stimuli channels were filled with fresh cell culture medium with 10% FBS to produce nutrient gradient along the narrow migration channels. After 6 hours of incubation, cells across the entire chip were fluorescently labeled by loading 5 μM SYTO 24 (Invitrogen) to both the sample and stimulation channels and incubate for 15 min. Migration behaviour of the cells were inspected under a fluorescence microscope. Cells that entered the migration channels or reached the stimuli channels were considered as deformed and migrated cells. The number of migrated cells was counted and compared for the deformability and migration potential of the cells from different samples. For testing the effect of rotenone on deformability and migration of the cells, rotenone at IC_20_ was added to the medium loaded onto the chip.

### Transwell cell invasion assay

To prepare the invasion assay chambers, 750 μL of 10% FBS-supplemented RPMI-1640 was added to each of the 24 wells of the Transwell plate (Corning), and 100 μL of Matrigel (Corning) was added to the cell culture inserts (8 μm). The plate was incubated in a humidified incubator at 37 °C with 5% CO_2_ for an hour. 500 μL of PC-3M NTC or *SLIT2* KO cells in serum-free RPMI-1640 were added to the Matrigel-coated inserts at 1×10^5^ cells/mL. Based on the conditions tested, media in the wells was either with or without 10% FBS and 1μg/mL secreted N-SLIT2 (R&D Systems, 8616-SL-050). The cells were incubated in the Matrigel-coated inserts for overnight in the incubator. The next day, medium was removed from the inserts and wells. The non-migrated cells on the upper side of the membrane was removed using a cotton swab and washed away with PBS. Cells remained on the membrane was fixed by submerging the membranes in the wells filled with cold methanol for 20 min and air-dried for 30 min. The membranes were then submerged in 750 μL 0.1 % crystal violet (Sigma) in wells for 30 min at room temperature to stain the cells before washed thoroughly with distilled water. After air-drying, membranes were submerged in 750 μL of 10 % acetic acid (VWR) with shaking until completely dissolving the stain. Optical density of the dissolved stain in each well was determined at 590 nm using a microplate reader.

### Oncology proteome antibody array

Oncology antibody array (R&D Systems) was performed following the manufacturer’s instructions. Briefly, NTC or *SLIT2* KO cell lysates were diluted and incubated overnight with the nitrocellulose membranes that had been pre-spotted in duplicates with antibodies targeting 84 oncology-related proteins. The membranes were washed before incubated with a cocktail of biotinylated detection antibodies, followed by addition of Streptavidin-HRP. The amount of protein bound at each capture spot was detected using chemiluminescent substrate and the signal was captured using a ChemiDoc imaging system (Bio-Rad). Intensity of the capture spots were quantified using ImageJ.

### Immunoblot analysis

Whole cell extracts were prepared with RIPA buffer (ThermoFisher) supplemented with protease inhibitor cocktail (Sigma). Proteins were separated on 4-15 % precast protein gels (Bio-Rad) and transferred onto PVDF membranes. Primary antibodies used were anti-SLIT2 (Abcam, ab134166), anti-Vimentin (Abcam, ab92547), anti-N-Cadherin (BD Bioscience, 610920), anti-E-cadherin (R&D Systems, AF648), and anti-Snail (NEB, 3879S). Secondary antibodies were purchased from CST. Blots were visualized with chemiluminescent substrate (Thermo Scientific) using a ChemiDoc imaging system. Intensity of the protein bands were quantified using ImageJ.

### RNA-sequencing and data analysis

RNA was extracted using RNeasy Plus Kit (Qiagen) following manufacturer’s instructions and submitted to PMGC for cDNA library preparation and RNA sequencing using Illumina NovaSeq 6000. RNA-sequencing data was processed for sgRNA representation using A*STAR procedure. Figures were generated using the normalized read counts in R and RStudio (R project, Revolution Analytics), Metascape, and the Gene Set Enrichment Analysis tools (Broad Institute).

### Cell viability assay

NTC or *SLIT2* KO PC-3M cells were seeded on 96-well plates in triplicates at a density of 2×10^4^ cells per well. The next day, rotenone was titrated into the 96-well plate at two-fold serial dilution with concentration ranging from 0-10 μM. Viable cell population was determined 48 hr later using Cell Counting Kit-8 (Dojindo) following the manufacturer’s instructions.

### Clinical and patient data analysis

RNAseq and clinical data was retrieved from portals of TCGA, the Human Protein Atlas, GEPIA2, and COSMIC. *SLIT2* expression median of 2.45 FPKM was used as the threshold to define low- or high-expression of *SLIT2* in prostate cancer patients (PRAD). *SLIT2* expression levels in tumor and normal tissues were retrieved for 31 cancer types to assess fold change. Information on the loss-of-function mutations in the top screen hits occurred in cancer patients was retrieved from Cancer Mutation Census (CMC) at COSMIC to compared with the level of loss-of-function mutations in well-known TSGs and OGs ^28^.

## Notes

### Competing Interest Statement

The authors have declared no competing interest.

