## Extended Data for "Genome-wide in vivo screen of circulating tumor cells identifies SLIT2 as a regulator of metastasis"

*Contents*

**Extended Data Fig.1:** Microfluidic CTC isolation from mice whole blood and PCR amplification of barcoded sgRNA region from mice genome. **page 2**

**Extended Data Fig. 2:** Pairwise comparison of gRNA abundance of the transduced cell pool before transplant, primary tumor, and CTC samples. **page 3**

**Extended Data Fig. 3:** Top enriched sgRNAs identified in primary tumors of the CTC screen targeted tumor suppressor genes. **page 4**

**Extended Data Fig. 4:** Fold change of *SLIT2* expression in tumor tissues compared to normal tissues across various cancer types. **page 5**

**Extended Data Fig. 5:** *SLIT2* KO xenograft mouse model results. **page 6**

**Extended Data Fig. 6:** Quantitation of cells in 3D spheroids formed from the non-targeted control and the *SLIT2* KO cells. **page 7**

**Extended Data Fig. 7:** Microfluidic device used to test cell deformability and migration. **page 8**

**Extended Data Fig. 8:** 2D cell growth of control and *SLIT2* KO cells. **page 9**

**Extended Data Fig. 9:** Proteomic profiling of the *SLIT2* KO cells. **page 10**

**Extended Data Fig. 10:** Surface EpCAM level of the control and the *SLIT2* KO cells. **page 11**

**Extended Data Fig. 11:** ATP6 and multiple components of complex I were upregulated in *SLIT2* KO cells. **page 12**

**Extended Data Fig. 12:** Knockout of *SLIT2* in LNCaP and PC3. **page 13**
**Extended Data Fig. 13:** Rotenone sensitivity in other *SLIT2* KO prostate cancer cells. **page 14**
**Extended Data Fig. 14:** Invasiveness of other *SLIT2* KO prostate cancer cells. **page 15**
**Supplementary Table 1:** Top enriched gene KOs in primary tumors. **page 16**
**Captions for SI_1, SI_2, and SI_3** **page 17**

**Supplementary Information for this manuscript includes the following:**

SI_1, SI_2, SI_3 (.xlsx)

**Extended Data Fig. 1: Microfluidic CTC isolation from mice whole blood and PCR amplification of barcoded sgRNA region from mice genome.**

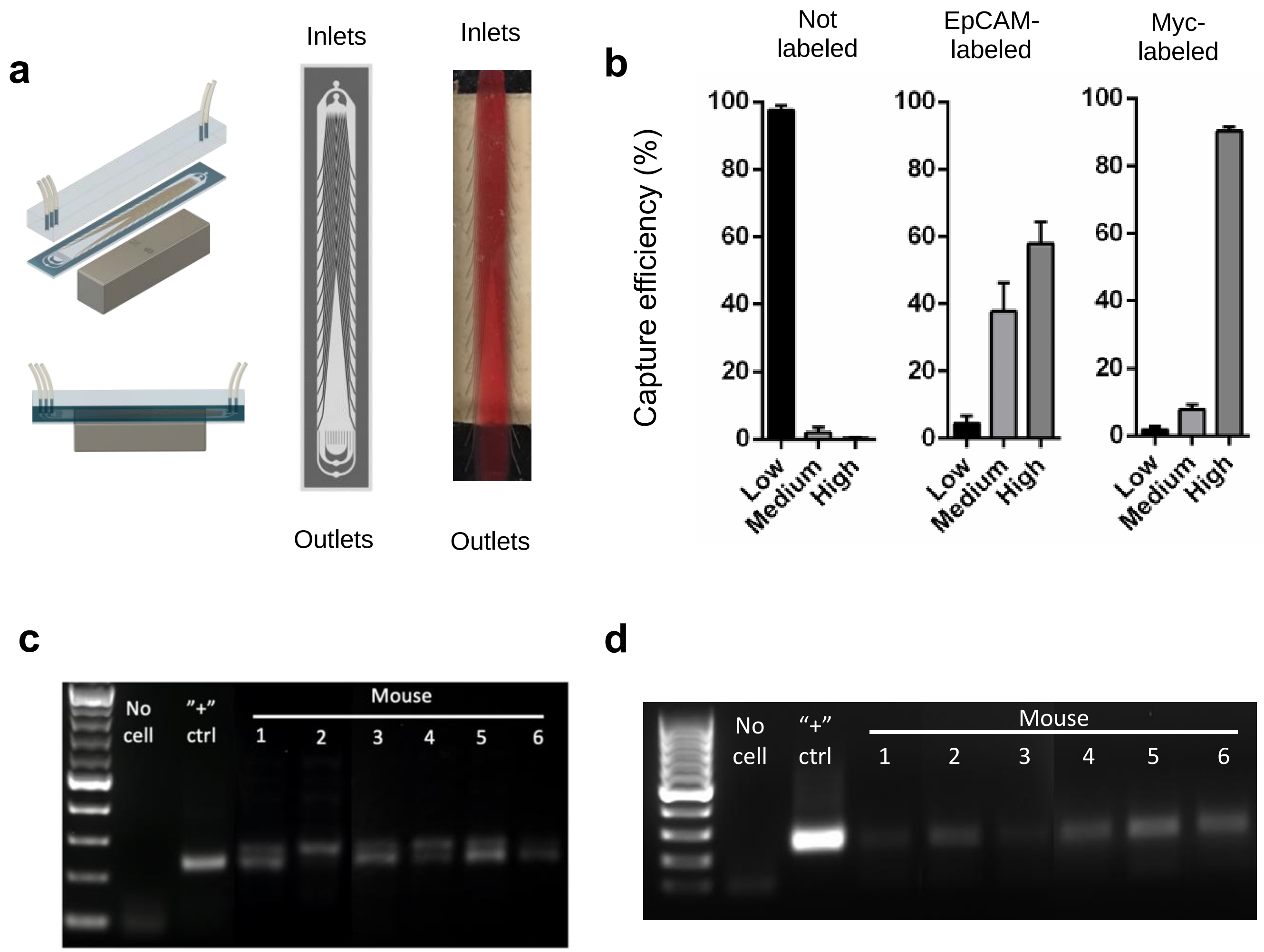


**a,** Images of Prism chip. Top left: schematic of Prism chip (3D view). Bottom left: schematic of Prism chip (side view). Middle: schematic of Prism chip (top view). Right: photo of a Prism chip processing mouse whole blood sample (top view). **b,** Bar graph demonstrating performance of Prism chip in terms of percent of cells collected in low-, medium-, and high-expression outlets. From left to right: PC-3M cells unlabeled, labeled with EpCAM magnetic microbeads, and labeled with biotin-conjugated Myc antibody + anti-biotin magnetic microbeads. **c,** Agarose gel image showing the PCR products of the barcoded sgRNA regions amplified from the CTC genome 3 weeks after the immunocompromised mice were transplanted with the TKOv3-transduced PC-3M cells. **d,** Agarose gel image showing the PCR products of the barcoded sgRNA regions amplified from the CTC genome 3 weeks after the immunocompromised mice were transplanted with the sub-pool-transduced PC-3M cells.

**Extended Data Fig. 2: Pairwise comparison of sgRNA abundance of the transduced cell pool before transplant, primary tumor, and CTC samples.**

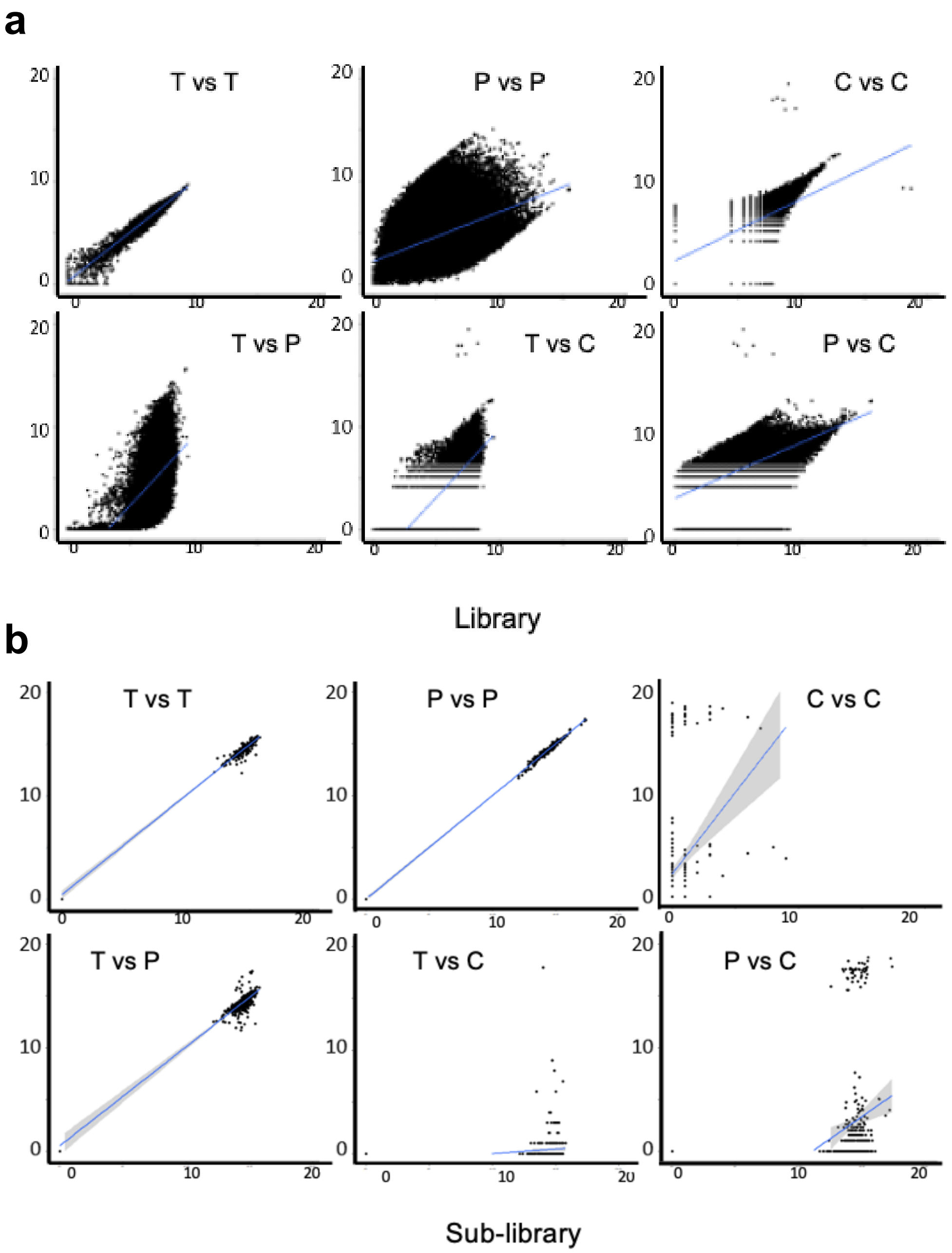


**a,** Pearson correlation of the normalized $\log_2$ sgRNA counts from the individual samples of the genome-wide CRISPR KO screen. T: replicate of the transduced cell pool before transplant. P: randomly selected replicate of primary tumor samples. C: randomly selected replicate of CTC samples. **b,** Pearson correlation of the normalized $\log_2$ sgRNA counts from the individual samples of the sub-pool screen. T: replicate of the transduced cell pool before transplant. P: randomly selected replicate of primary tumor samples. C: randomly selected replicate of CTC samples.

**Extended Data Fig. 3: Top enriched sgRNAs identified in primary tumors of the CTC screen targeted tumor suppressor genes.**

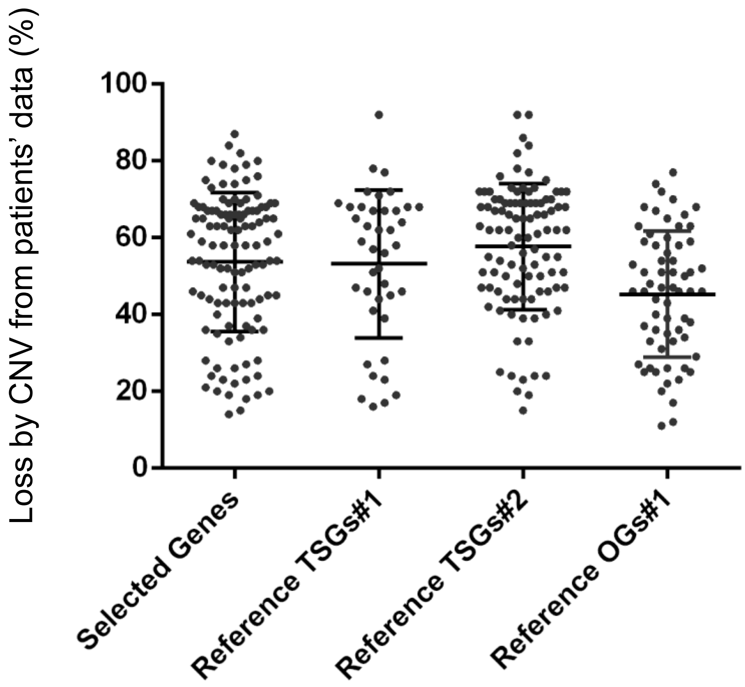


Dot plot comparing the percentage of loss-of-function of each top selected gene from the genome-wide CTC CRISPR screen (Selected Genes) and the percentage of loss-of function of the tumor suppressor genes (Reference TSGs) and oncogenes (Reference OGs) selected from published literature and COSMIC database. Gene loss-of-function and copy number variation (CNV) data of cancer patients is available on COSMIC.

**Extended Data Fig. 4: Fold change of *SLIT2* expression in tumor tissues compared to normal tissues across various cancer types.**

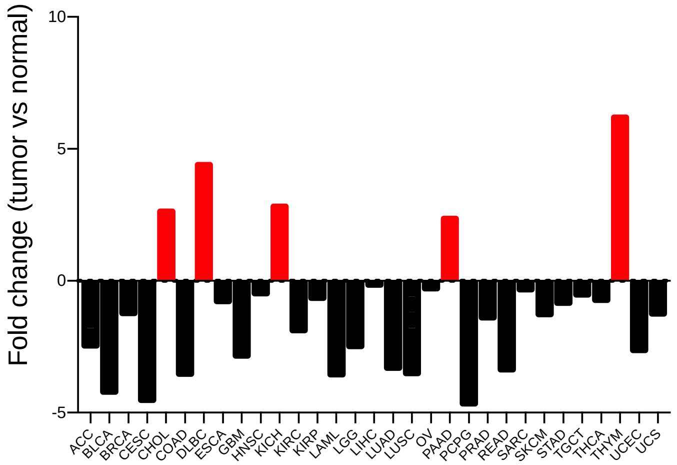


| | |
|---|---|
| **ACC** | adrenocortical carcinoma |
| **BLCA** | bladder urothelial carcinoma |
| **BRCA** | breast invasive carcinoma |
| **CESC** | cervical squamous cell carcinoma and endocervical adenocarcinoma |
| **CHOL** | cholangiocarcinoma |
| **COAD** | colon adenocarcinoma |
| **DLBC** | lymphoid neoplasm diffuse large B-cell lymphoma |
| **ESCA** | esophageal carcinoma |
| **GBM** | glioblastoma |
| **HNSC** | head and neck squamous cell carcinoma |
| **KICH** | kidney chromophobe |
| **KIRC** | kidney renal clear cell carcinoma |
| **KIRP** | kidney renal papillary cell carcinoma |
| **LAML** | acute myeloid leukemia |
| **LGG** | low grade gliomas |
| **LIHC** | liver hepatocellular carcinoma |
| **LUAD** | lung adenocarcinoma |
| **LUSC** | lung squamous cell carcinoma |
| **OV** | ovarian serous cystadenocarcinoma |
| **PAAD** | pancreatic adenocarcinoma |
| **PCPG** | pheochromocytoma and paraganglioma |
| **PRAD** | prostate adenocarcinoma |
| **READ** | rectum adenocarcinoma |
| **SARC** | sarcoma |
| **SKCM** | skin cutaneous melanoma |
| **STAD** | stomach adenocarcinoma |
| **TGCT** | testiculat germ cell tumors |
| **THCA** | thyroid carcinoma |
| **THYM** | thymoma |
| **UCEC** | uterine corpus endometrial carcinoma |
| **UCS** | uterine carcinosarcoma |

*SLIT2* expression levels (median) in tumor and normal tissues of patients from 31 types of cancers were compared using RNAseq data collected from TCGA. Black bars indicate cancers with reduced expression of *SLIT2* in tumor compared to normal tissues, and the red bars indicate cancers with enhanced expression of *SLIT2* in tumor compared to normal tissues. Table on the right shows the full names of the abbreviations of the 31 cancer types.

**Extended Data Fig. 5: *SLIT2* KO xenograft mouse model results.**

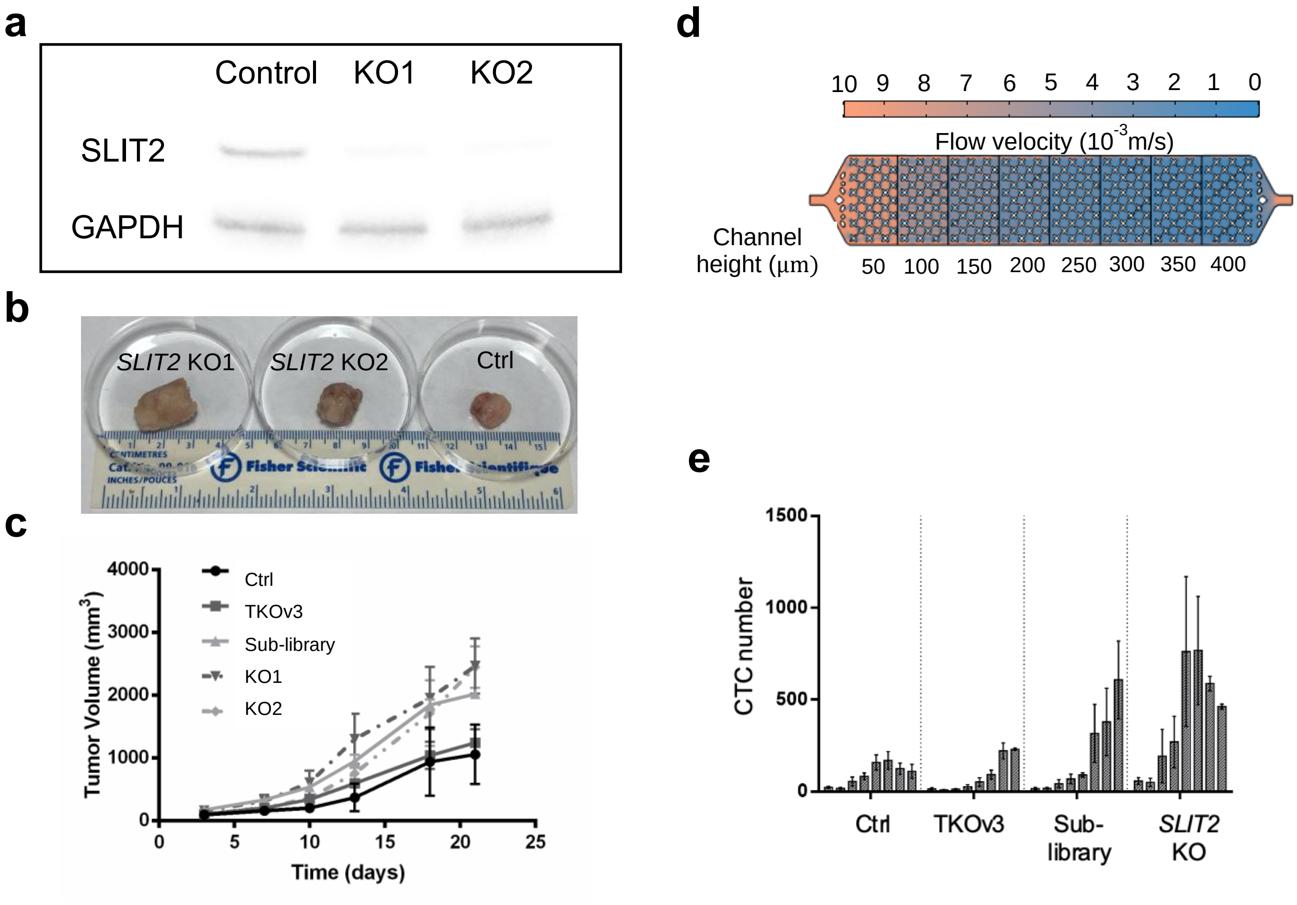


**a,** Immunoblot image showing successful knockout of *SLIT2* in PC-3M cells. KO1 and KO2 are knockout cell lines generated using two different sgRNAs. **b,** Representative photo of the primary tumors collected 3 weeks after the immunocompromised mice being transplanted with control or *SLIT2* KO PC-3M cells. **c,** Tumor growth curves comparing tumor size resulted from transplant of control, *SLIT2* KO, and CRISPR library-transduced cells (n = 3). **d,** Schematic of the 8-zone CTC capture and counting device, featuring the “x”-shaped cell-capture pockets. The 8 capture zones have the same width but increasing height from top (inlet) to the bottom (outlet), with the color scheme demonstrating the flow velocity. **e,** Bar graph of the CTC number counted from the immunocompromised mice transplanted with control, *SLIT2* KO, and CRISPR library-transduced cells (n = 3). This graph shows detailed CTC number counted from each zone of the 8-zone CTC-capture device for each transplant group. Error bars indicate SD.

**Extended Data Fig. 6: Quantitation of cells in 3D spheroids formed from the non-targeted control and the *SLIT2* KO cells.**

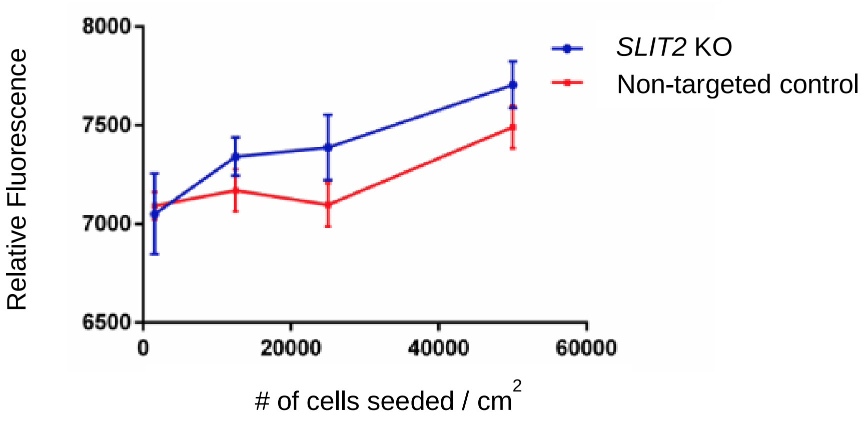


Non-targeted control and *SLIT2* KO PC-3M cells were plated at various cell density to grow 3D spheroids. Cell numbers of the 3D spheroids in each well was determined based on the amount of ATP present using the CellTiter-Glo 3D cell viability assay.

**Extended Data Fig. 7: Microfluidic device used to test cell deformability and migration.**

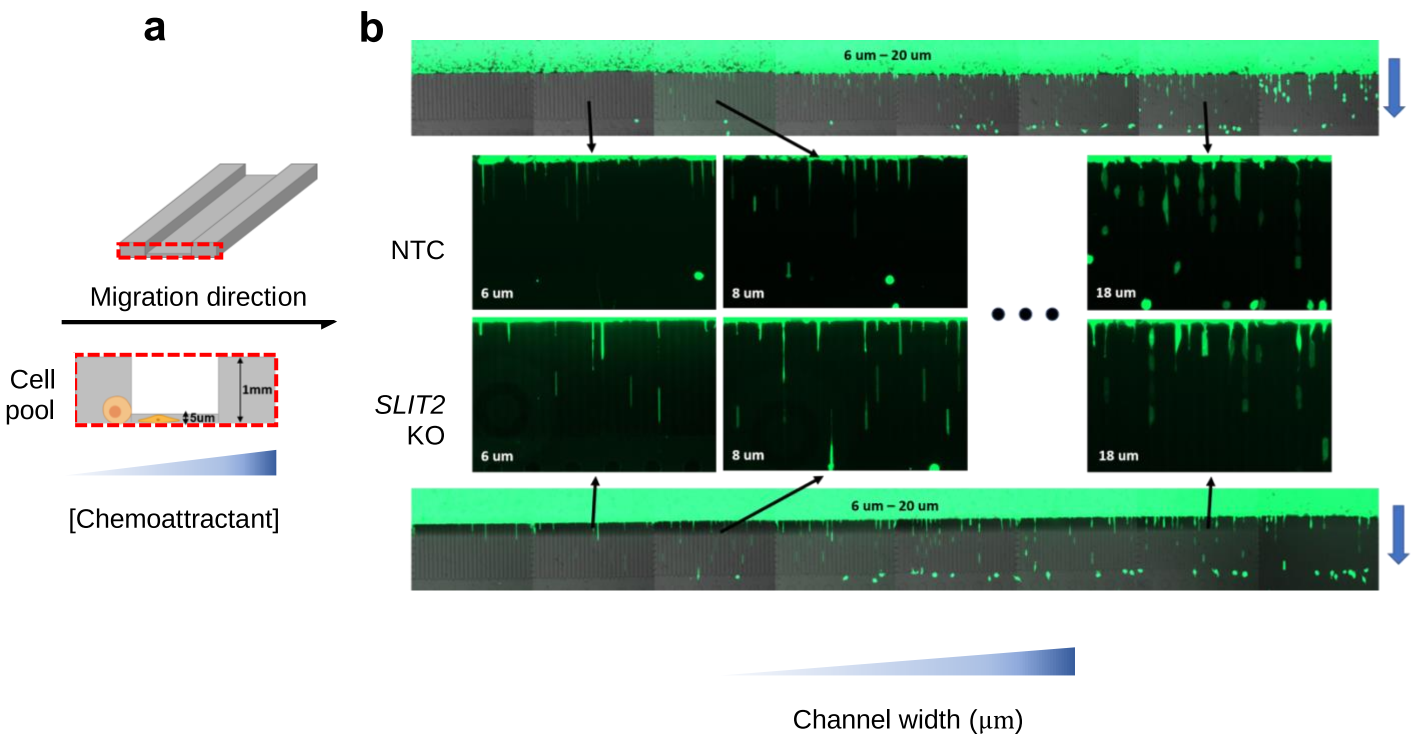


**a,** Schematic of the migration device challenging the cells of their ability to deform and travel through the channels towards the chemoattractant. **b,** Representative microscopic images of the migration microfluidic device 24 hours after being loaded with the non-target control or *SLIT2* KO cells. The overall device images sandwich the zoom-in images of the sections with the same channel height, but varying width as indicated by the blue wedge at the bottom. The blue arrows on the right indicate the direction of cell migration.

**Extended Data Fig. 8: 2D cell growth of the non-targeted control and the *SLIT2* KO cells.**

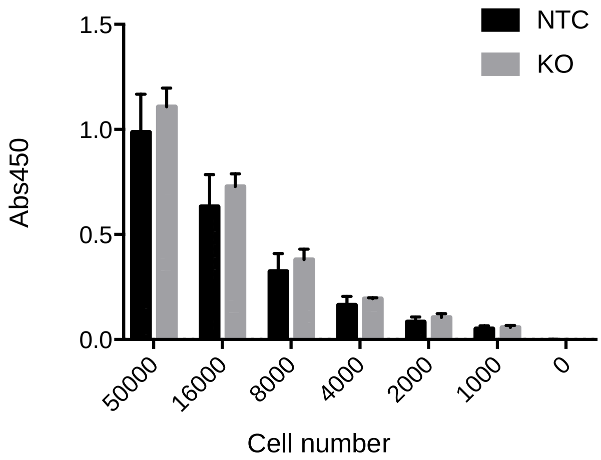


Non-targeted control and *SLIT2* KO PC-3M cells were plated at various cell density on a 96 well plate and cultured for 3 days (n = 3 per cell line per cell density). At the end of culturing, cell numbers were determined using CCK-8 assays.

**Extended Data Fig. 9: Proteomic profiling of the *SLIT2* KO cells.**

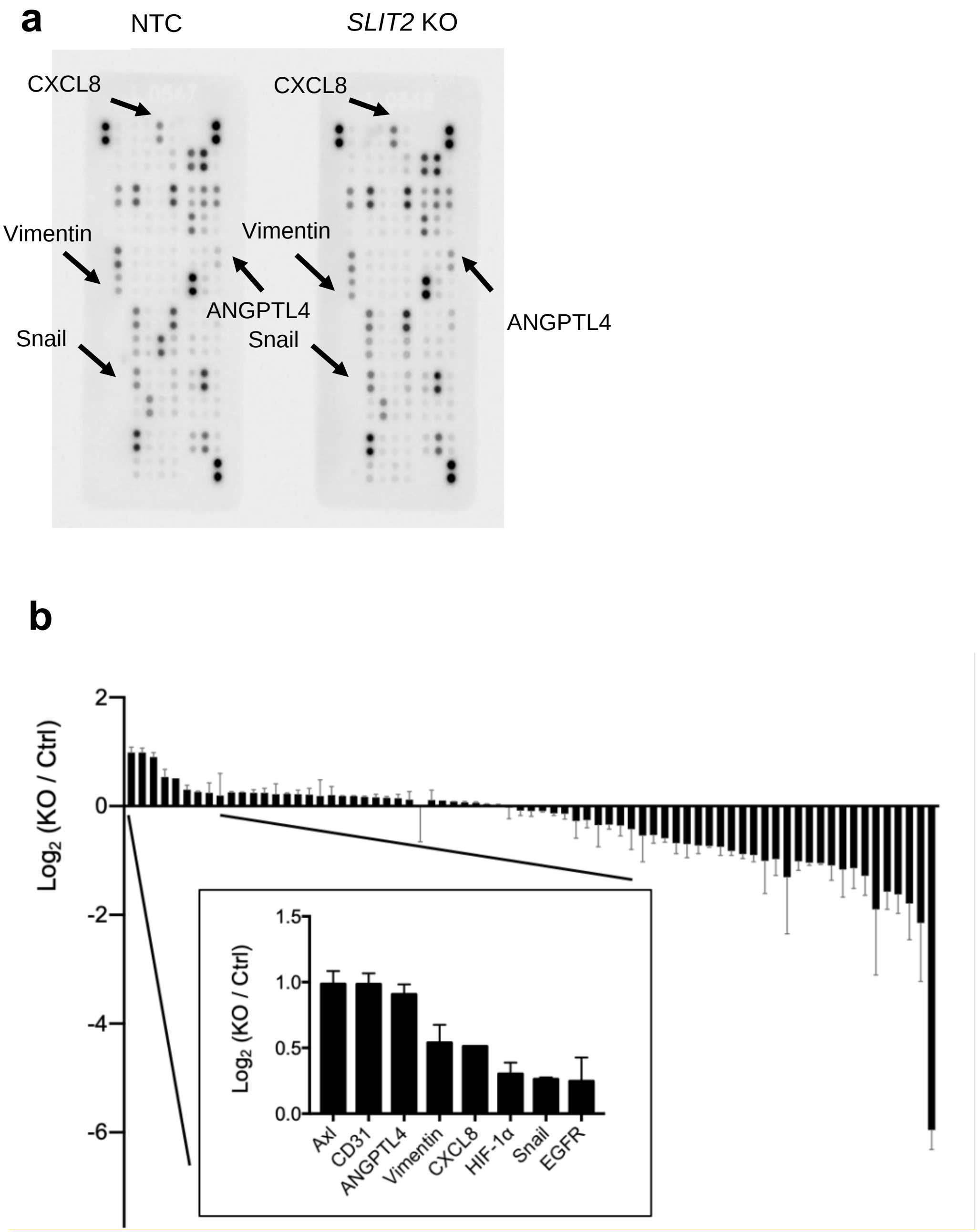


**a,** Blot images of the oncology antibody array performed with the non-targeted control and the *SLIT2* KO cells. Each dot represents detection of one type of protein, with 2 replicates. The most up-regulated proteins in the *SLIT2* KO cells found in the oncology antibody array were labeled with arrows. **b,** Bar graph showing the ranking of tested proteins based on the fold change of expression in *SLIT2* KO cells compared to the non-targeted control in the oncology antibody array. Expression levels were compared based on the dot intensities in (**a**) quantified using Image J.

**Extended Data Fig. 10: Surface EpCAM level of the non-targeted control and the *SLIT2* KO cells.**

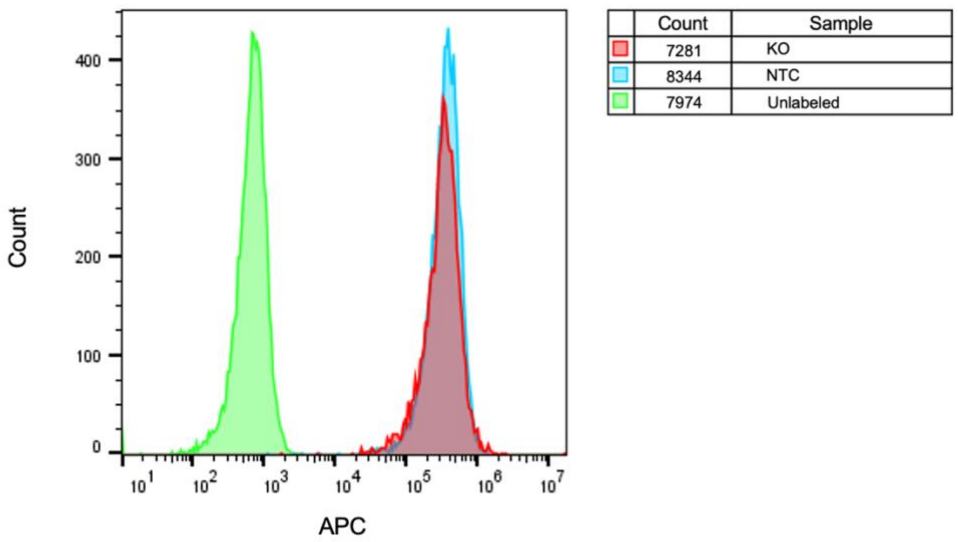


Non-targeted control and *SLIT2* KO PC-3M cells were labeled with EpCAM magnetic microbeads followed by APC-conjugated anti-beads antibody. Data was processed using FlowJo.

**Extended Data Fig. 11: ATP6 and multiple components of complex I were upregulated in *SLIT2* KO cells.**

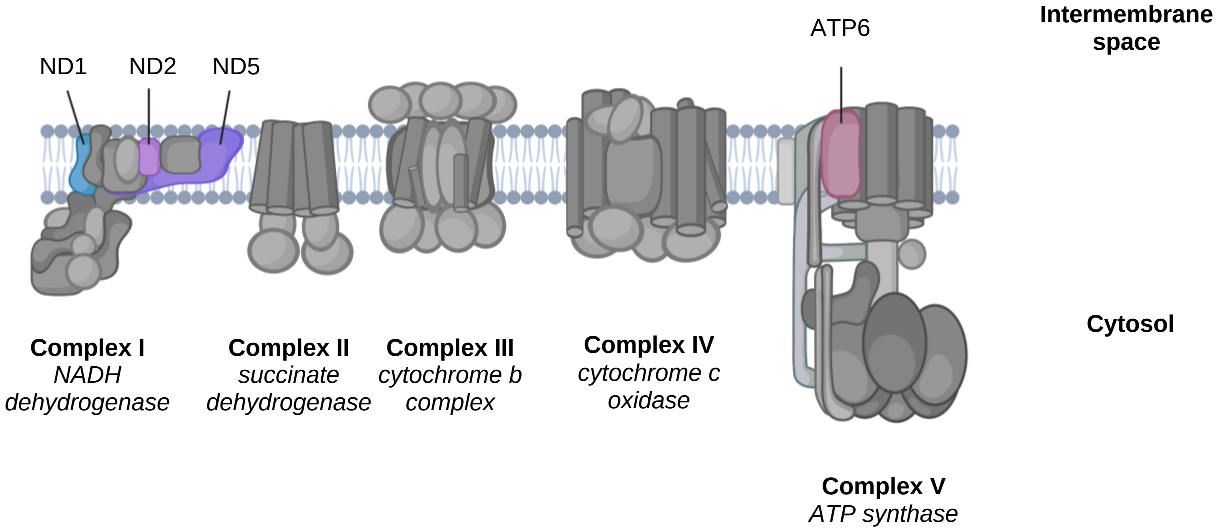


Schematic of the electron transport chain in mitochondria, highlighting the components that correspond to the most up-regulated group of genes identified from RNA-seq of the *SLIT2* KO cells.

**Extended Data Fig. 12: Knockout of *SLIT2* in LNCaP and PC3 using CRISPR technology.**

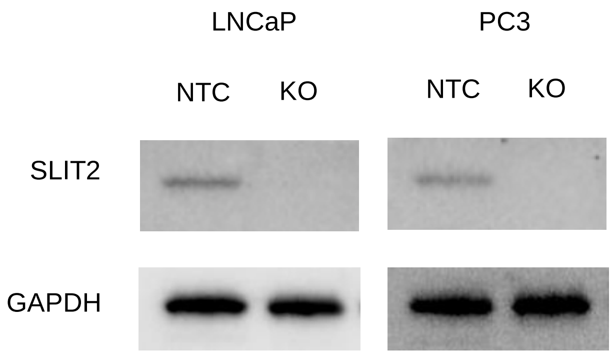


Immunoblot images showing successful knockout of *SLIT2* in LNCaP and PC3 cells. GAPDH was blotted as house-keeping gene.

**Extended Data Fig. 13: The effect of *SLIT2* on rotenone sensitivity of other prostate cell lines.**

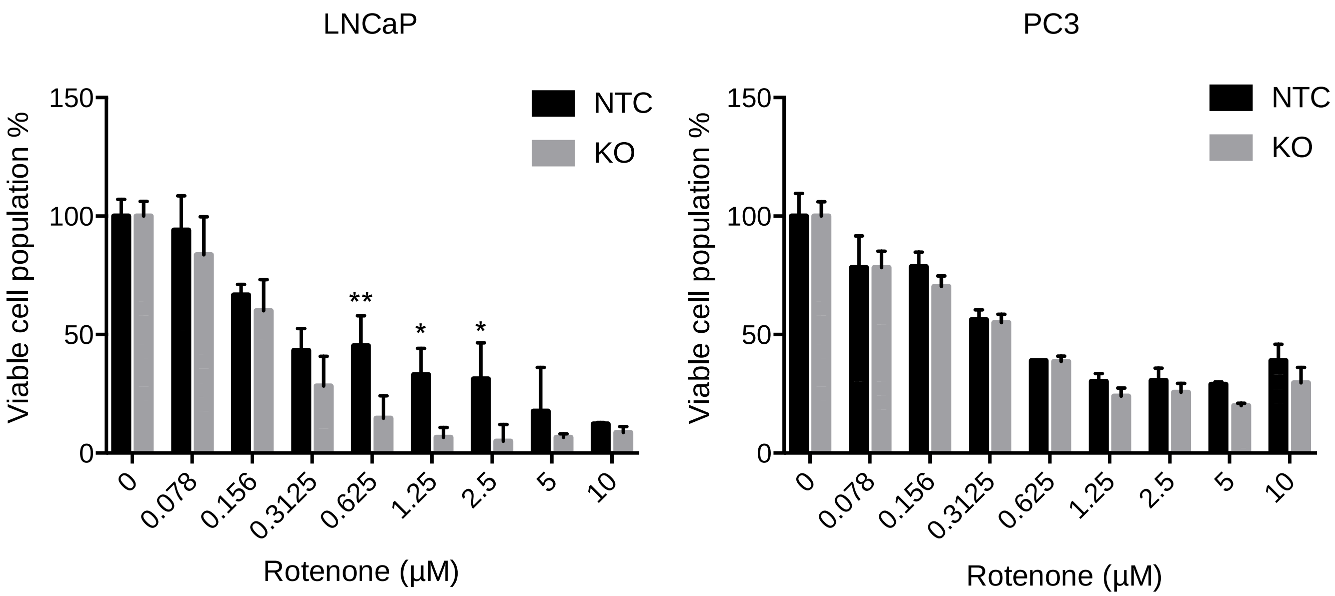


Bar graphs showing viability of the *SLIT2* KO compared to NTC cells (LNCaP on the left, PC3 on the right) under titration of rotenone (n = 3 for each group). Error bars indicate SD. P values were calculated by two-way ANOVA. ****P<0.0001; ***P<0.001; **P<0.01; *P<0.05.

**Extended Data Fig. 14: The effect of *SLIT2* on the invasiveness of other prostate cell lines.**

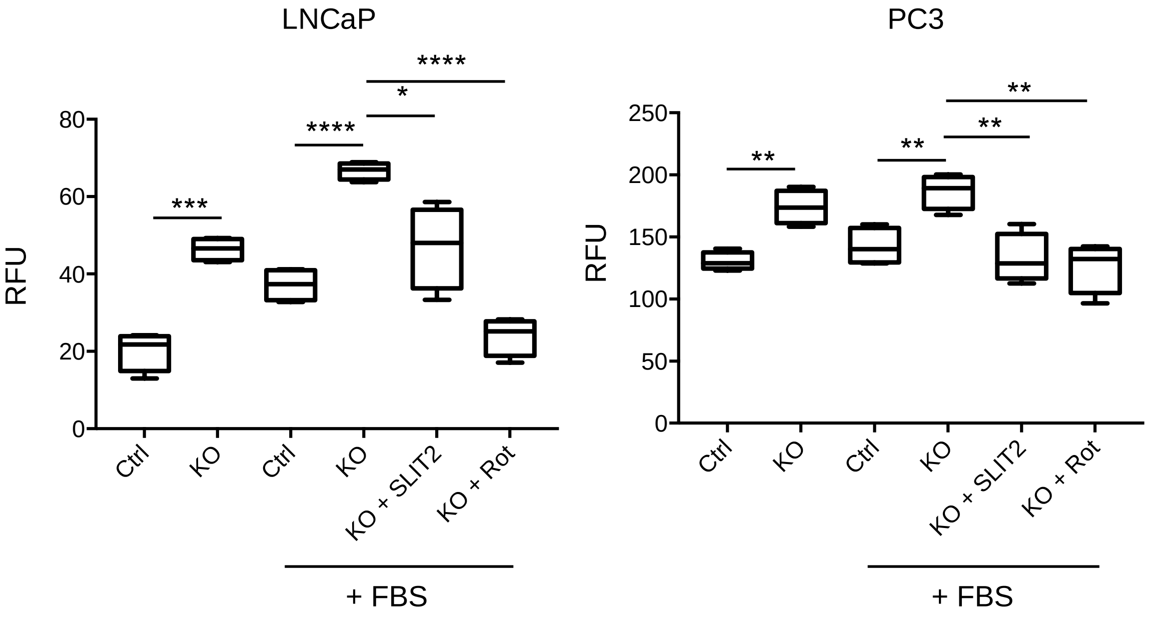


Box plots comparing the number of NTC versus *SLIT2* KO cells (LNCaP on the left, PC3 on the right) invaded across the transwell membrane with or without the presence of the nutrient, N-SLIT2 protein, rotenone in the culture media (n = 4 for each group). Rotenone at $IC_{20}$ was used. For LNCaP, 2 μg/mL N-SLIT2 was used; for PC3, 4 μg/mL N-SLIT2 was used. P values were calculated by two-tailed unpaired t-test. ****P<0.0001; ***P<0.001; **P<0.01; *P<0.05.

| GENE | FDR |
| --- | --- |
| TSC1 | 0.000104 |
| PSG3 | 0.000525 |
| PROSER2 | 0.00209 |
| RFPL2 | 0.0049 |
| LLGL1 | 0.00733 |
| DACH1 | 0.0191 |
| FAXC | 0.0191 |
| TMUB1 | 0.0203 |
| USP42 | 0.0203 |
| CACHD1 | 0.0222 |
| PHF19 | 0.0232 |
| GTF2I | 0.0278 |
| SLC25A13 | 0.0363 |
| REEP5 | 0.038 |
| SLIT2 | 0.0454 |
| ARSJ | 0.0454 |
| LMAN1 | 0.0454 |
| ZRSR1 | 0.0454 |
| PIGL | 0.0461 |

**Supplementary Table 1: Top enriched gene KOs in primary tumors.**Top ranked gene KOs in the primary tumors compared to the initial cell pool based on false discovery rate (FDR < 0.05).

SI_1. (Separate file)
CTC Genome-wide CRISPR KO Screen Results

SI_2. (Separate file)
CTC Sub-library CRISPR KO Screen Results

SI_3. (Separate file)
Genes and percent loss in Extended Data Fig. 3
